# Coupled yawing and orbital rotation in kinesin motility revealed by polarization measurement of gold nanorods

**DOI:** 10.1101/2021.11.07.467662

**Authors:** Mitsuhiro Sugawa, Yohei Maruyama, Masahiko Yamagishi, Robert A. Cross, Junichiro Yajima

## Abstract

Kinesin motor domains generate impulses of force and movement that have both translational and rotational components, raising the question of how the rotational component contributes to motor functions. We used a new assay in which kinesin-coated gold nanorods (kinesin-GNRs) move on suspended microtubules, for two plus-end-directed and weakly-processive kinesins: single-headed KIF1A, dimeric ZEN-4. Polarization of the light scattered by two types of kinesin-GNRs periodically oscillated as they orbited the microtubule along a left-handed helical trajectory. Our analyses revealed that each kinesin-GNR unidirectionally rotates about its yaw axis as it translocates, and that the period of this yaw-axis rotation corresponds to two periods of its left-handed helical orbit around the microtubule axis. Theoretical analyses suggest that the yaw-axis rotation enhances biased lateral displacement of the kinesin team. Our study reveals biaxial rotation as a new mode of motility in kinesin teams that may help the team to sidestep roadblocks.

## Introduction

Kinesins are adenosine 5’-triphosphate (ATP)-driven and microtubule-based molecular motors with multiple mechanical functions, including cargo transport along microtubules, the regulation of microtubule dynamics, and the sliding of bundled microtubules(1–3). Recent advances in techniques for 3D particle tracking and simultaneous measurements of force and torque have revealed that kinesins have translational and rotational degrees of freedom on microtubules, such that their motilities consist of both helical(4–8) and rotational(9) motions. For example, single-molecule kinesin-1 dimers unidirectionally rotate their stalk domain as they move processively via a hand-over-hand mechanism, especially at high ATP concentration(9). Mutagenic elongation of the neck linker of kinesin-1 dimers increases the probability of protofilament switching during stepping along the microtubule(10), so that the neck-linker mutant dimers retain processivity but switch protofilaments more readily, driving left-handed helical motion of multiple motor-coated beads around the axis of the microtubule (6). These studies suggest that the protofilament-tracking ability of wild type kinesin-1 dimers may derive from their rotary hand-over-hand mechanism(9) and their neck stability(6, 8). Wild-type kinesin-2 and −8, which have longer neck-linker domains than kinesin-1, and kinesin-6, which has an atypical neck region, do not track protofilaments. All these kinesins move towards microtubule plus ends along left-handed helical trajectories, as shown in a bead assay(6, 11), a microtubule surface-gliding assay(8, 11–13) and a single-molecule tracking assay(8). Furthermore, teams of non-processive dimeric kinesin-14 Ncd(14–16), which moves towards microtubule minus ends, as well as teams of truncated single-headed kinesins-1(17, 18), −3(19), −5(4, 20) and −6(11) motors have all been shown to drive corkscrewing motions of microtubules in surface-gliding assays. The handedness of these motions is consistent. Whilst plus-end directed kinesins move along a left-handed helix, minus-ended kinesins move along a right-handed helix. These helical motilities imply that the stroke (the unitary impulse of force and movement generated by the motor domains) has an off-axis component, driven either by conformational changes in the motor domain or by the directionally biased selection of microtubule binding sites (or potentially both)(6, 8, 11, 15, 17, 19). The underlying mechanisms and functional relevance of the torque component remain elusive.

To gain further insight, we have developed a novel motility assay in which gold nanorods (GNRs), coated with a team of kinesin molecules, move unhindered along suspended microtubules. Our assay uses light scattered by the kinesin-coated GNRs to track their position, and the polarization of the scattered light to report their orientation. We performed the GNR motility assay on three plus-end-directed kinesins: single-headed KIF1A, dimeric ZEN-4.

## Results

### GNR imaging for particle tracking and polarization measurements

We developed a motility assay based on tracking of kinesin-coated GNRs. Focusing of light scattered by the GNRs allows positional tracking, whilst the ratio of the polarization of this scattered light in orthogonal directions reports the orientation of the GNR as it moves over the microtubule surface (Fig. 1A and Materials and Methods). GNR images were obtained by laser dark-field imaging using highly inclined illumination with nearly total internal reflection angle(21, 22) via a perforated dichroic mirror(23). The position of each GNR spot was measured in three dimensions using a cylindrical lens(24) (Fig. S1). Polarization measurements of GNRs were performed using a combination of the polarized incident laser beam and a polarizing beam displacer, which two orthogonal polarization signals were projected on the camera, allowing us to measure 0° to 180° GNR orientations in the sample plane (Fig. 1B and Materials and Methods). Kinesin constructs were prepared with a biotinylated peptide (Avi-tag) fused to their C-terminus. Biotinylated GNRs (40 nm × 68 nm) were coated in streptavidin and biotinylated kinesin molecules were attached to this streptavidin layer (Materials and Methods). We used the GNR with relative low aspect ratio to detect various rotations of kinesin coated GNR (kinesin-GNR) on the microtubule. A GNR with higher aspect ratio may inhibit rotations of a kinesin-GNR, since the parallel longitudinal axis alignment of the kinesin-GNR and the microtubule is most stable in terms of the number of kinesin molecule that can attach to the microtubule. Maximum capacity of kinesin molecules attached to the GNR used in this study is roughly estimated to be about 100 molecules (surface area of GNR/occupied area of kinesin ≈ 1.1 × 10^4^ nm^2^/10^2^ nm^2^) and about one fourth of the attached kinesin molecules could generate driving force and torque on the microtubule. Microtubules were suspended on microbeads (0.5 μm) via the antigen–antibody system for β-tubulin, so that kinesin-coated GNRs were free to orbit the microtubules between bead-attachment points (Fig. 1C). Control experiments established that the Z-displacement of GNR in the range of ±200 nm had negligible effect on the GNR polarization measurement in our experimental system (Fig. S2). Using the GNR motility assay, we performed a comparative analysis of two plus-end-directed kinesins, which have distinctive properties of processivity and oligomeric state: biased-diffusional motile and single-headed human kinesin-3 (KIF1A)(25) and weakly processive and dimeric *Caenorhabditis elegans* kinesin-6 (ZEN-4)(26).

**Figure 1.**
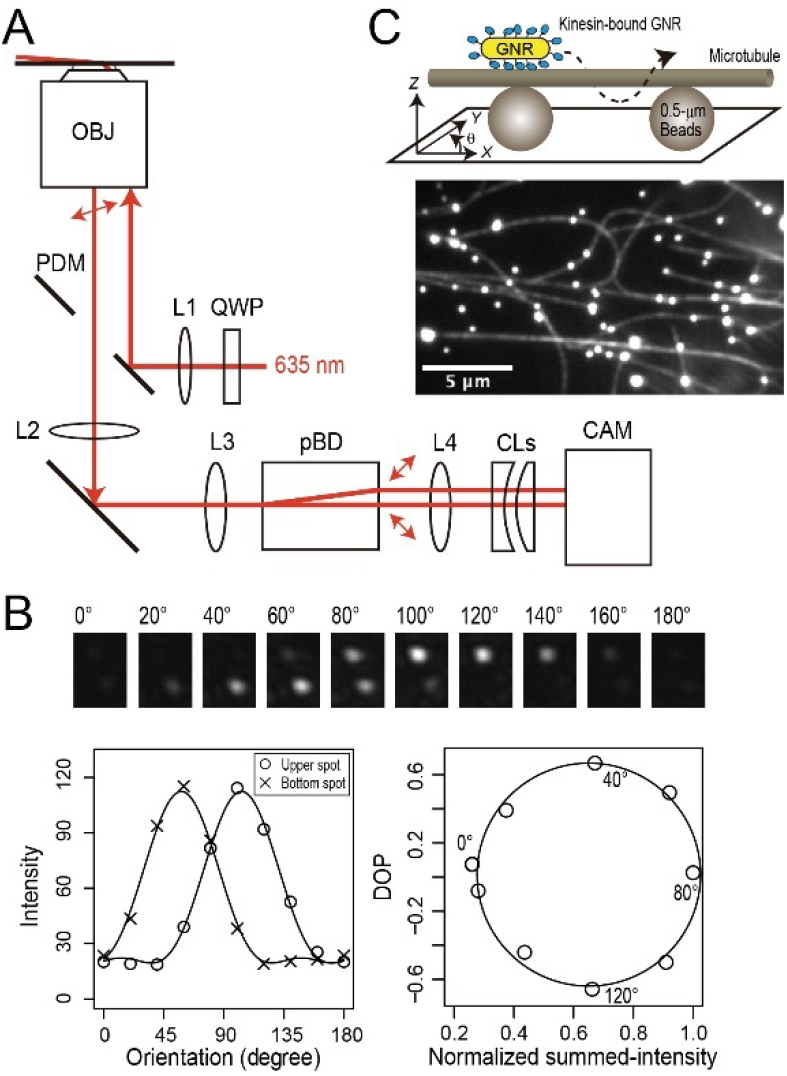
GNR motility assay. (**A**) Schematic drawing of the optics for the gold nanorod (GNR) motility assay. OBJ, objective lens; PDM, perforated dichroic mirror; QWP, quadruple wave plate; pBD, polarizing beam displacer; CL, cylindrical lens; CAM, CMOS camera; L, lens. (**B**) Calibration for polarization measurements. (upper) Series of pairs of scattered-light spots of microbeads (0.1 μm) modulated by a polarizer, which was located in front of L3, for emulating GNR spots with various orientations in the plane. (lower left) Intensity profiles of a pair of scattered-light spots (*I*_bottom_, *I*_upper_) shown in the upper images. The plots of *I*_bottom_ and *I*_upper_ were fitted by intensity functions of the GNR orientation (Materials and Methods). (lower right) Degree of polarization (DOP), (*I*_bottom_ – *I*_upper_)/(*I*_bottom_ + *I*_upper_) as a function of the normalized summed-intensity of two scattered-light spots, (*I*_bottom_ + *I*_upper_). (**C**) (top) Schematic drawing of the GNR motility assay in which a kinesin-coated GNR moves along a microtubule suspended with fluorescent microbeads (0.5 μm) attached to a glass substrate. (bottom) Fluorescence image of Cy5-labelled microtubules suspended on fluorescent microbeads. The bright spots are the fluorescent microbeads.

### GNR motility assay for single-headed KIF1A

We performed the GNR motility assay for single-headed KIF1A (1-366 amino acid residues (AAs))(25) at a saturating ATP concentration (2 mM ATP). GNRs carried multiple KIF1A molecules, allowing GNRs to move processively along microtubules. We found that the light-scattering intensities of each pair of KIF1A-coated GNR (KIF1A-GNR) spots, corresponding to orthogonal polarization signals from a single GNR, periodically oscillated in intensity during helical movement, with an increase in each signal corresponding to a decrease in the other (13 particles out of 62 tested KIF1A-GNRs) (Fig. 2A and Movie S1). The degree of polarization (DOP) as a function of the summed-intensity of the two GNR spots depicted an elliptical trajectory, indicating continuous rotations of the polarization (Fig. 2B). The *X-Y-Z*-trajectory of the KIF1A-GNR showed steady left-handed helical motion around the microtubule long axis (Fig. 2C). Figure 2D shows that the time trajectories of the *X*- and *Y*-displacements, the DOP and summed-intensity, and the polarization angle as viewed from above the glass substrate. The time trajectory of the polarization angle exhibited continuous counterclockwise rotations during the left-handed helical motion of the KIF1A-GNR around the microtubule. The *Y*-displacement and the polarization signals of the KIF1A-GNR exhibited strong cross-correlation and periodicity (Fig. 2E), indicating that the GNR polarization were synchronized with the helical motion on the microtubule. We also observed clockwise and oscillatory rotations of the GNR polarization, synchronized with the helical motion of KIF1A-GNRs (Fig. 3A and 3B). The translational velocity (parallel to the microtubule axis) was 0.51 ± 0.12 μm s^−1^ (mean ± standard deviation (SD), *n* = 13 GNRs), which is nearly equal to the microtubule sliding velocity of 0.49 μm s^−1^ in the surface-gliding assay (Fig. S3). The pitch of the helical motion measured in the *X-Y*-trajectory was 0.69 ± 0.36 μm (mean ± SD, *n* = 33 cycles in 13 KIF1A-GNRs), which is slightly shorter than that of the corkscrewing of microtubules on a surface of KIF1A of 0.89 ± 0.21 μm (mean ± SD, *n* = 35 cycles in 5 microtubules) (*P* ≪ 0.01, Welch’s *t* test). The displacement in the microtubule axis during one periodic change in the GNR polarization signals (hereafter referred to as polarization pitch) was 0.69 ± 0.37 μm (mean ± SD, *n* = 33 cycles in 13 KIF1A-GNRs), which was almost equal to that of the helix pitch (Fig. 3C). We note that the measured helix pitch is a combination of the helix pitch exerted by motors on the straight microtubule lattice and any supertwist pitch of the microtubule, with the supertwist depending on the number of the protofilaments. However, when the measured pitch is less than 1 μm, as it is here, the supertwist of the microtubule has negligible influence on the measured pitch (Fig. S4 and Materials and Methods). We therefore report the uncorrected helix pitch unless otherwise mentioned.

**Figure 2.**
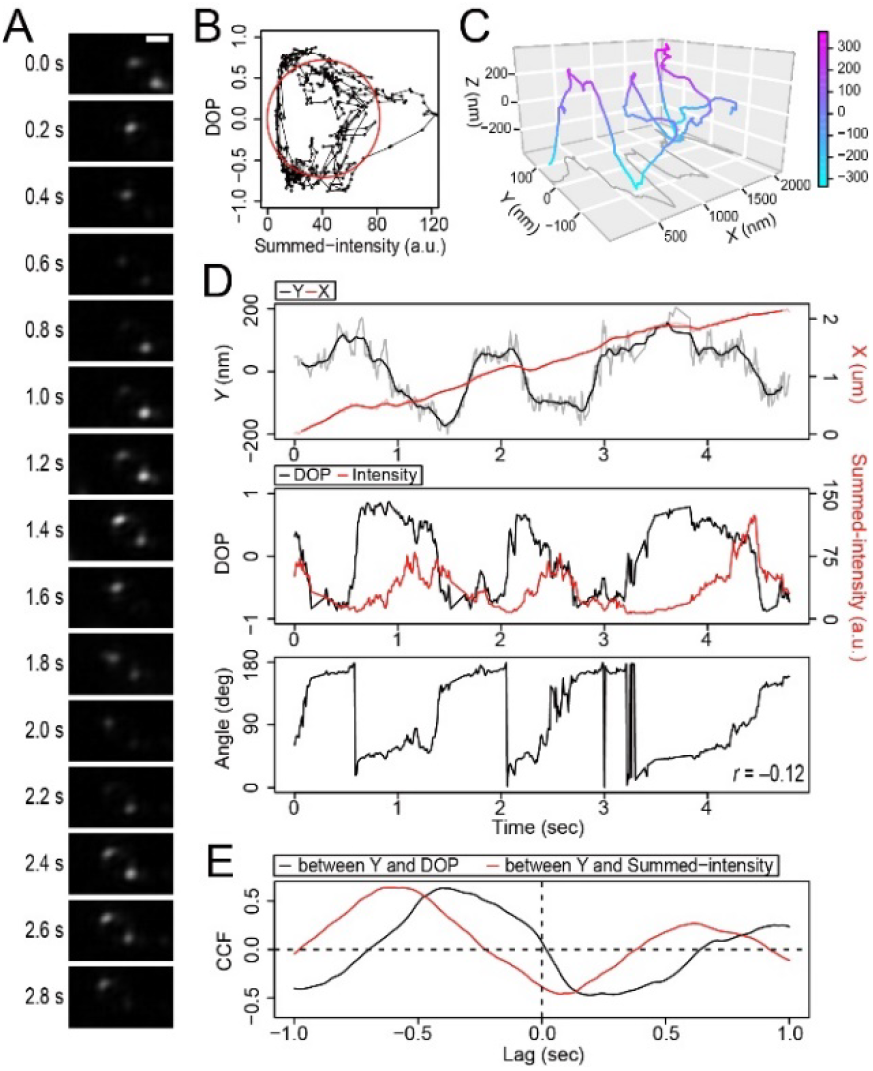
Typical data of the GNR motility assay for KIF1A. (**A**) Montage of scattered-light images of one KIF1A-coated GNR (KIF1A-GNR) moving around a microtubule every 0.2 s in the presence of 2 mM ATP. The frame rate was 100 frames s^−1^. Scale bar, 2 μm. (**B**) Degree of polarization (DOP) as a function of the summed-intensity of the two KIF1A-GNR spots shown in (A). The red line represents an ellipse fit. (**C**) 3D trajectory of the KIF1A-GNR shown in (D) through a moving-average filter of 15 frames. (**D**) The time trajectories of the *X*- and *Y*-displacements (upper), the DOP and summed-intensity of the two spots (middle), and the estimated angle (bottom) of KIF1A-GNR shown in (A). The value of *r* is Pearson’s correlation coefficient between the *Y*-displacement and the angle. (**E**) Cross-correlation functions (CCFs) between the *Y*-displacement and the polarization signals.

**Figure 3.**
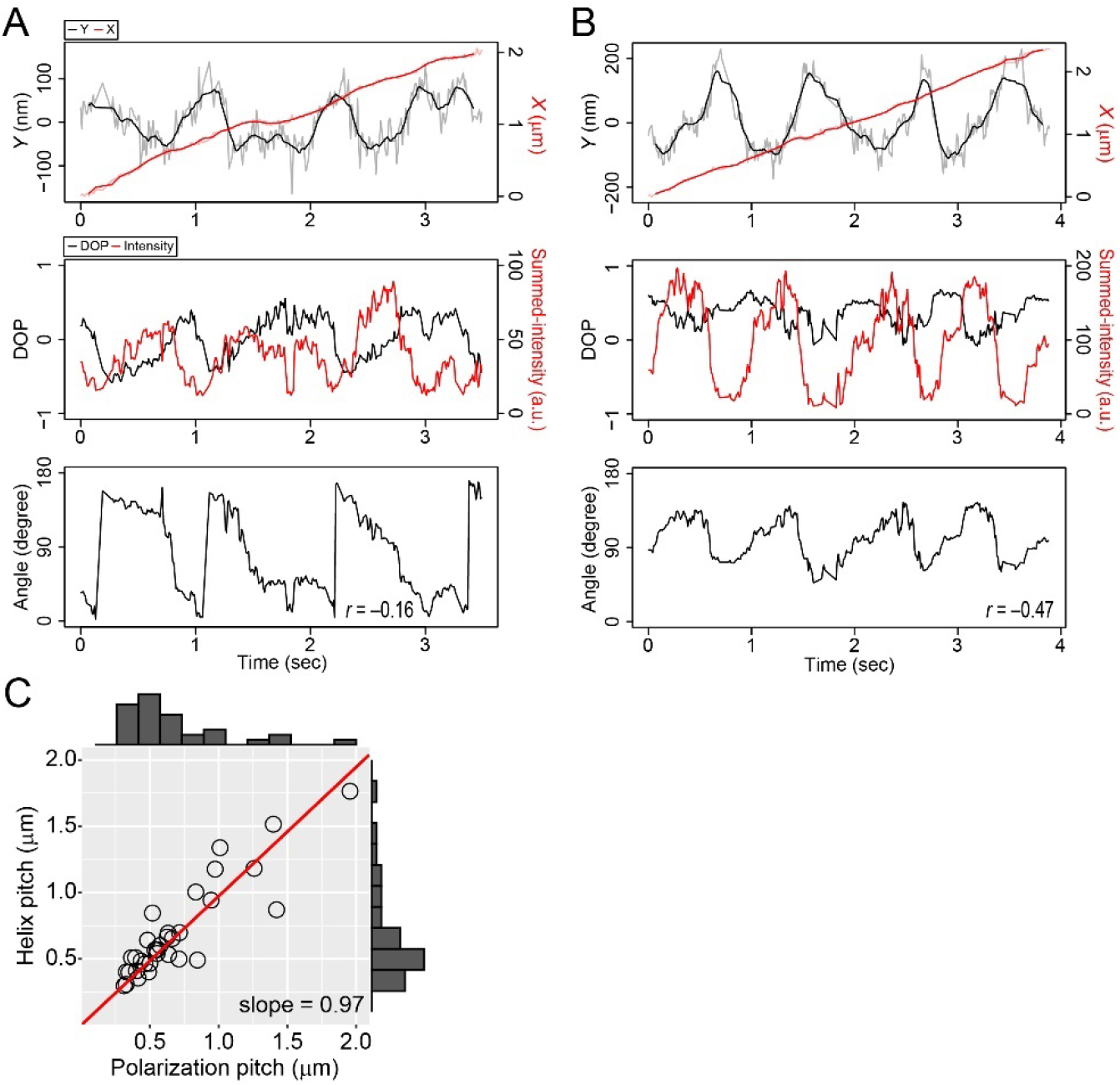
Coupled polarization rotation and helical motion of KIF1A-GNRs. (**A, B**) The time trajectories of the *X*- and *Y*-displacements (upper), the degree of polarization (DOP) and summed-intensities of the two spots (middle), and the estimated angles (bottom) of KIF1A-GNRs. The time trajectories of the polarization in the image plane show clockwise rotation in (A) and oscillatory rotation in (B). The value of *r* is Pearson’s correlation coefficient between the *Y*-displacement and the angle. (**C**) Distribution of the helix and polarization pitches of KIF1A-GNRs. The helix and polarization pitches are the displacements during one periodic change in the *Y*-displacement and the polarization, respectively. The red line represents a linear fit (*y* = *ax*) of which the slope (*a*) is 0.97 (*R*^2^ = 0.95).

### Helical and yaw-axis rotations of kinesin-coated GNRs

The GNR motility assays of KIF1A revealed that the polarization of KIF1A-GNRs rotated periodically whilst moving along a helical trajectory toward microtubule plus ends. As a control to check whether rotation of the GNR polarization is coupled to helical orbiting of the supertwist of the microtubules or not, we studied GNRs coated with multiple kinesin-1 coated GNRs, which faithfully track protofilaments(6) rather than side-stepping. Polarization angle of the kinesin-1 coated-GNRs did not exhibit unidirectional rotation along a long-pitch helical path in our observation (Fig. S5). When the orientation of a kinesin-GNR relative to the microtubule axis is constant during helical motion, unidirectional rotations of the GNR polarization are never observed (Fig. 4A and Movie S2). Therefore, we hypothesized that unidirectional rotations of the GNR polarization are generated by rotations in three axes (rolling, yawing, pitching) of the kinesin-GNR during helical motion. We note that considering symmetry of gold nanorods and geometries of our polarization measurement, pitching and yawing of the GNR cannot be distinguished from each other in our polarization measurement. However, as shown below, the experimental results can be well explained by the combination of rolling and yawing in the kinesin-GNR. We constructed a model, in which a kinesin-coated GNR unidirectionally rotates about its yaw and roll axes during the short-pitch helical motion around the microtubule long axis (Movie S3). At first, time trajectories of the kinesin-coated GNR are given by

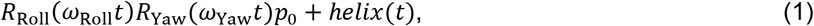

where *R*_Roll_(*ω*_Roll_*t*) is rolling matrix with the angular velocity *ω*_Roll_, *R*_Yaw_(*ω*_Yaw_*t*) is yawing matrix with the angular velocity *ω*_Yaw_, and *helix*(*t*) is a helical displacement as functions of time *t*, and *p*_0_ is an initial position of a kinesin-GNR with an initial phase angle *φ*. Note that the initial phase angle *φ* is defined by the relative angle to the microtubule axis when the kinesin-coated GNR is at the top position of its helical trajectory. The polarization angle measured in this study is the orientation of the GNR projected in the *X-Y*-plane. Then, time trajectories of the GNR angle in the *X-Y*-plane are given by

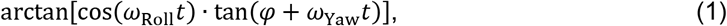

**Figure 4.**
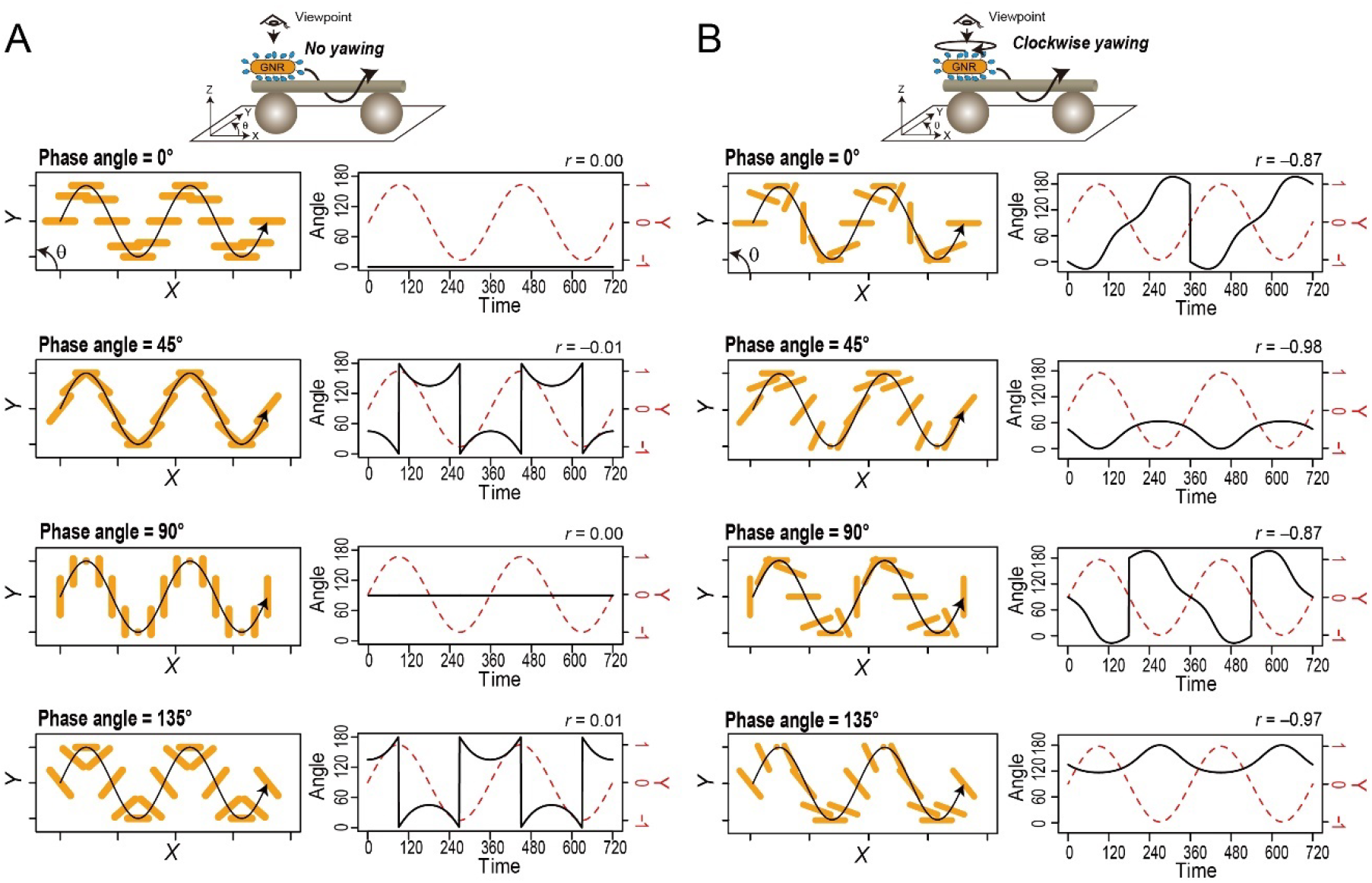
Interpretation of the observed polarization rotations and translational movement of the kinesin-coated gold nanorods (GNRs). (**A, B**) (left) Montage of the projection images of a GNR in the *X-Y* plane, obtained by a model of a motor-coated GNR with no yawing (A) or 180° clockwise yawing during one period of helical motion (B). (right) Time trajectories of the *Y*-displacement and angle of a motor-coated GNR with different initial phase angles (*φ* = 0°, 45°, 90°, and 135°). The value of *r* is Pearson’s correlation coefficients between the *Y*-displacement and the angle. Trajectories of a motor-coated GNR in the case of counterclockwise yawing are shown in Fig. S6. See also Movies S2 and S4.

Results of KIF1A-GNRs were well explained when *ω*_Roll_ = ±2*ω*_Yaw_. This means that a KIF1A-coated GNR unidirectionally rotates 180° about its short (yaw) axis in one period of short-pitch helical motion around the microtubule long axis. The three types of rotations of the GNR polarization angle (counterclockwise, oscillatory, and clockwise rotations) can be reproduced by different initial phase angles (*φ*) of the model (Fig. 4B and Movie S4). For example, rotations of the GNR polarization are counterclockwise at *φ* = 0°, oscillatory at *φ* = 45° and 135°, and clockwise at *φ* = 90°. Since the initial phase angle *φ* is a random variable in the assay, our model is consistent with experimental results in which these three types of polarization rotations of KIF1A-GNRs were observed.

In this model, the GNR yawing direction is also a determinant of apparent rotational patterns of the GNR polarization in the sample plane. Pearson’s correlation coefficient between the Y displacement and the angle of a kinesin-GNR is negative when it yaws clockwise and positive when counterclockwise (Fig. 4B and Fig. S6). Estimated by the correlation coefficient between the *Y*-displacement and the polarization angle, 8 out of 13 KIF1A-GNRs exhibited clockwise yawing whilst 5 out of 13 KIF1A-GNRs counterclockwise yawing (see also Fig. 2D and Fig.3A and B). The directionalities of yawing of these kinesin-coated GNRs were not significantly different from an equal probability using binomial test (*P* > 0.5).

### GNR motility assay for dimeric ZEN-4

We performed the GNR motility assay for the weakly-processive dimeric kinesin, ZEN-4 (1-555 AAs)(26), again at a saturating ATP concentration (2 mM) (Materials and Methods). We again found that the polarization signals of the ZEN4-coated GNRs (ZEN4-GNR) periodically changed, correlated with the *Y*-displacements during steady forward movement (13 particles out of 49 tested ZEN4-GNRs) (Fig. 5A). The analyses revealed the counter-clockwise, clockwise, and oscillatory rotations of these ZEN4-GNRs. The forward velocity of the ZEN4-GNR was 0.39 ± 0.05 μm s^−1^ (mean ± SD, *n* = 13 ZEN4-GNRs), which is comparable to the velocity in the bead assay(11). The polarization pitch of the ZEN4-GNR was 0.67 ± 0.20 μm (mean ± SD, *n* = 34 cycles in 13 ZEN4-GNRs) in the microtubule axis, which is equal to the helix pitch of 0.67 ± 0.21 μm (mean ± SD, *n* = 34 cycles in 13 ZEN4-GNRs) (Fig. 5B). Thus, whilst ZEN4-GNRs moved more slowly than KIF1A-GNRs, the polarization rotation of ZEN4-GNRs was again phase-locked to their helical motion, as for KIF1A-GNRs. These results indicate that ZEN4-GNRs also unidirectionally rotates 180° about its short (yaw) axis in one period of short-pitch helical motion around the microtubule long axis. Estimated by the correlation coefficient between the *Y*-displacement and the polarization angle, 5 out of 13 ZEN4-GNRs exhibited clockwise yawing, whilst 8 out of 13 ZEN4-GNRs counterclockwise yawing. The directionalities of yawing of these ZEN4-GNRs were not significantly different from an equal probability using binomial test (*P* > 0.5).

**Figure 5.**
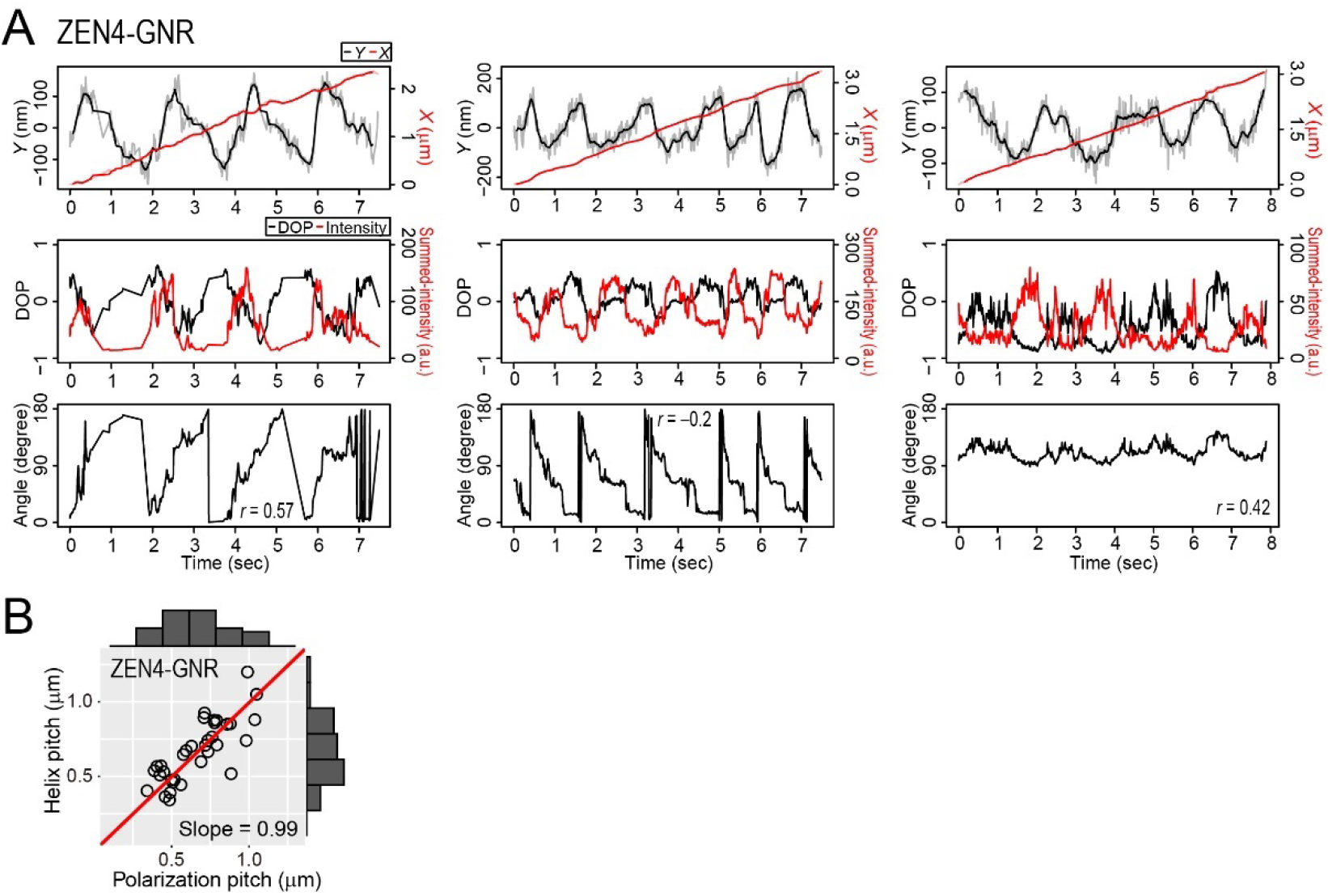
GNR motility assay for ZEN-4. (**A**) Typical time trajectories of the *X*- and *Y*-displacements (upper), the degree of polarization (DOP) and summed-intensity of the two spots (middle), and the estimated angles (bottom) of ZEN-4-coated gold nanorods (ZEN4-GNRs). The three trajectories represent counterclockwise (left), clockwise (middle), and oscillatory (right) rotations of the GNR polarization in the sample plane. The value of *r* is Pearson’s correlation coefficient between the *Y*-displacement and the angle. (**B**) Distribution of the helix and polarization pitches of ZEN4-GNRs. The red lines represent linear fits (*y* = *ax*) of which the slopes (*a*) are 0.99 (*R*^2^ = 0.97).

### Unidirectional rotation of microtubule on single ZEN4-GNR

Our proposed model, in which kinesin-GNRs rotate around their yaw axes as they move along and around the microtubule long axis, predicts that immobilized kinesin-GNRs might drive rotation of an overlying short microtubule about both its long and short axes. To test the rotational motility of the kinesin-GNRs during translation, we performed the surface-gliding assay for ZEN4-GNRs, in which ZEN4-GNR particles were sparsely attached on the glass surface (3-8 particles per 100 μm^2^) (Materials and Methods). We found that the short microtubules unidirectionally rotated on ZEN4-GNRs while only their ends were in contact with the GNRs (Fig. 6 and Movie S6). In the geometry of observation images, we observed both counterclockwise rotation (*n* = 4 microtubules) and clockwise rotation (*n* = 4 microtubules). These observations also support a model in which kinesin-coated GNRs moving along a microtubule are driven to undergo both helical and yaw-axis rotations.

**Figure 6.**
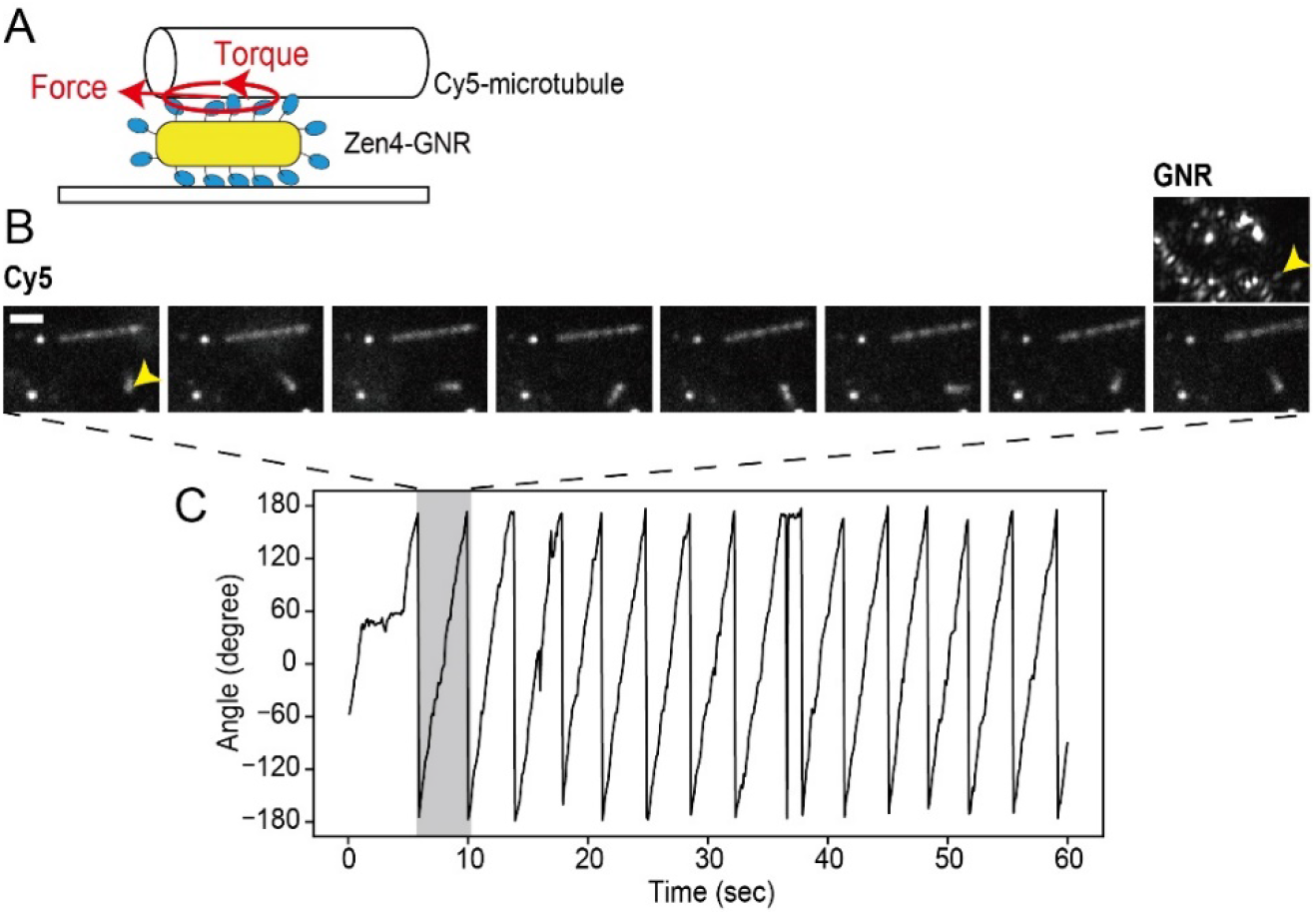
Surface-gliding assay for ZEN-4 coated gold nanorods (ZEN4-GNRs). (**A**) Schematic drawing of the surface-gliding assay, whereby the generation of force and torque rotate a Cy5-labelled microtubule (Cy5-microtubule) on the ZEN4-GNR. (**B**) Montage of fluorescence images of Cy5-microtubules every six frames (600 ms), showing that the short Cy5-microtubule continuously rotated (yellow arrowhead). Scale bar, 2 μm. GNR image of the same region-of-interest of the Cy5 images is also shown (right upper). (**C**) Time trajectory of the microtube orientation.

### Analysis of the kinesin-GNR motility by 2D ratchet mechanism

We found that the polarization and helix pitches were comparable among KIF1A- and ZEN4-GNRs (0.7-0.8 μm) (*P* > 0.05 for Welch’s *t* test), despite substantial differences in their forward velocities (*P* < 0.01 for adjusted pairwise comparisons using Welch’s *t* test with the Holm method). The helix pitch is given by *P_Helix_* = 2π*V*_X_/ω_Helix_, where *P_Helix_* is the pitch size of helix, *V*_X_ is the forward velocity, and ω_Helix_ is the angular velocity in the *Y-Z*-plane. We see that the forward velocities are different among KIF1A- and ZEN4-GNRs but the pitches are comparable, so that the ratios of *V*_X_ to ω_Helix_ are thus almost equal. This means that the angular velocity of helical motion is directly proportional to the forward velocity for two kinesin constructs, suggesting that the off-axis component is intrinsic to each power stroke. This relationship can be well modeled by a 2D noise-driven ratchet mechanism, as proposed by Mitra *et al*. for single-headed KIF1A(19). Based on this mechanism, we now consider how yawing of the kinesin-coated GNRs influences its motility. Since a kinesin-coated GNR rotated 180° about its yaw axis in one period of a helical trajectory with the helix pitch of about 700 nm, the GNR rotates 2° ≈ 180°/(700 nm/8 nm) with 8 nm of forward displacement, which is equivalent to the length of the tubulin dimer. The 2° rotation generates at most ±1 nm ≈ ±20~34 nm × tan(2°) of lateral displacements of the kinesin molecules bound on the GNR (40 nm in diameter and 68 nm in length) (Fig. 7A). Lateral displacement (*d*) of 1 nm is equivalent to 0.2 ≈ 1 nm/5.1 nm(29) in the asymmetry factor (*α*) toward the off-axis, which would be sufficient to influence the noise-driven ratchet mechanism (Fig. 7B). A recent discussion by Mitra et al. indicates that around 1 μm of a helix pitch is reproduced by the 2D Brownian ratchet model with a slight off-axis asymmetry (*α*_Y_ = 0.53) relative to the on-axis asymmetry (*α*_X_ = 0.27)(19).

**Figure 7.**
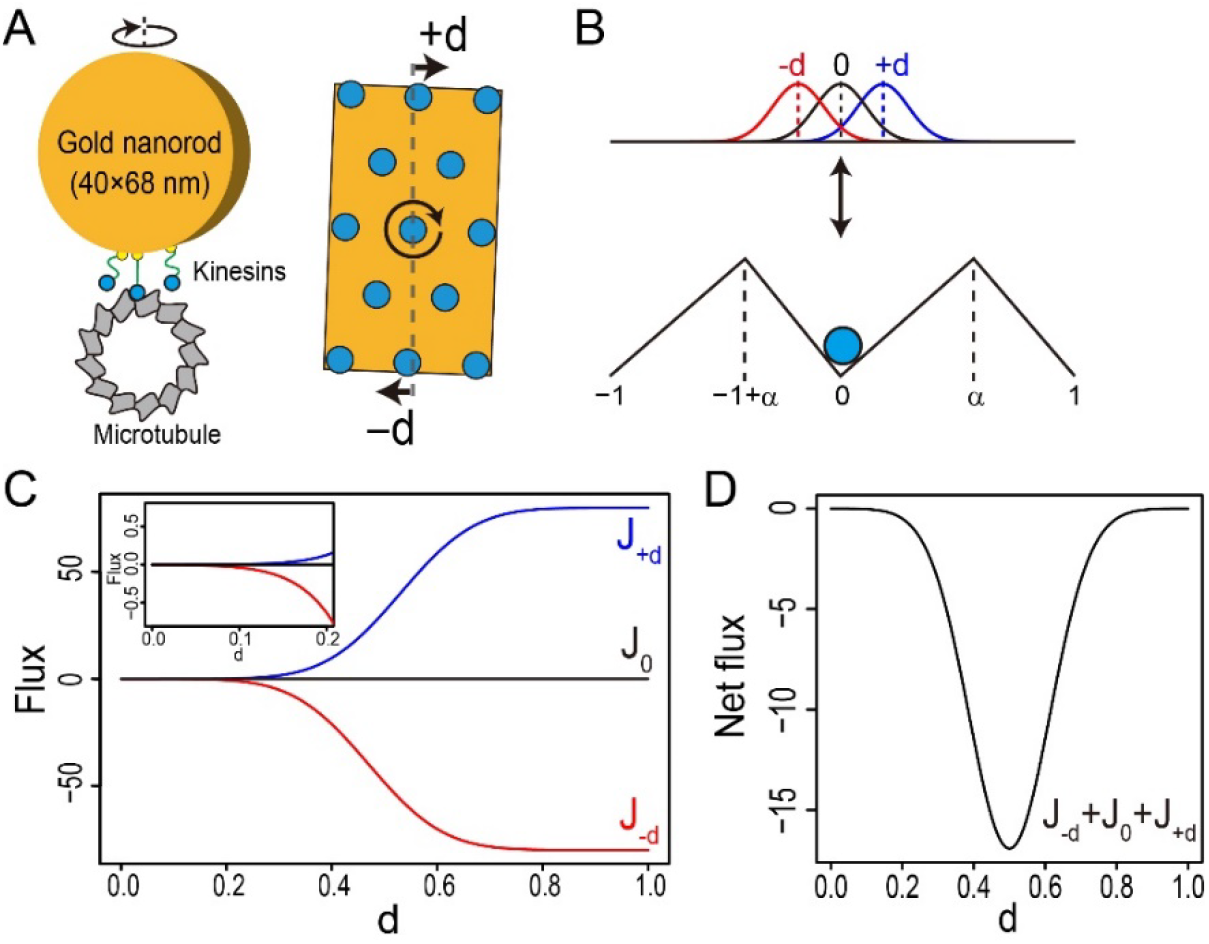
Mechanistic model for the clockwise rotation and helical motion of kinesin-coated gold nanorods (GNRs). (**A**) By rotation of the GNR, motors in the posterior and anterior positions relative to the rotation axis move laterally with displacement of ±*d*. (**B**) This situation can be modeled using the noise-driven ratchet model with asymmetry potentials (*a*) and three different starting positions of the particles (-*d*, 0, +*d*). (**C, D**) Fluxes of three particles at the starting positions of diffusion in the noise-driven ratchet with asymmetry potentials after 1/*γ* units of time (Materials and Methods). (**C**) Fluxes of three particles, *J_−d_*, *J*_0_, and *J_+d_* as a function of d with *α* = 0.53 and *γ* = 160 s^−1^. (**D**) Net flux, *J_−d_* + *J*_0_ + *J_+d_* as a function of *d*.

We evaluated the off-axis fluxes of three particles differing in their starting positions for diffusion (-*d*, 0, +*d*) in the noise-driven ratchet model with asymmetry potentials (*α*) after 1/*γ* units of time(30) (Materials and Methods):

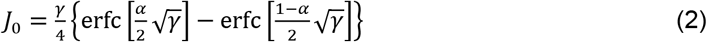

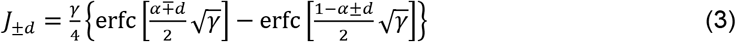

where *J*_0_ represents the flux of particles starting diffusion at 0 and *J_±d_* represents the flux of particles starting diffusion at ±*d*. In calculations with *α* = 0.53 and *γ* = 160 s^−1^ (Materials and Methods), the absolute values of *J_+d_* and *J_−d_* increased with d along a sigmoidal curve and the net flux, *J_−d_* + *J*_0_ + *J_+d_* described the convex downward function of which the value reached a minimum at *d* = 0.5 (Fig. 7C and 7D). Importantly, the slight displacement (*d* < 0.2) is sufficient to cause biased flux because |*J_−d_*| is much larger than |*J_+d_*| (Fig. 7C, inset). These suggest that unidirectional rotation of a motor-coated cargo enhances the sideward bindings of motor molecules toward the biased direction.

### Monte Carlo simulation of motor team with torque generation

To further explore the consequences of combined rotational and helical motion of kinesin-coated GNRs, we performed simple Monte Carlo simulations of the two-state noise-driven ratchet model. In this simulation, multiple particles, which are linked to a rigid body cargo with a spring, stochastically take biased forward and/or sideward steps on the 2D lattice and generate torque to twist the spring and then rotate the cargo when taking a step (Fig. 8A). Based on the force- and moment-balance equations, the cargo moves and rotates around the center of the joint points connecting the particles in the attached state (Materials and Methods). Additionally, the twist angle (Δ*θ*) of the cargo driven by the torque generated by the particles taking a step is given by Δ*θ* = (*N*_step_ *θ*_step_ + *b*_θ_)/*N*_on_, where *N*_step_ is the number of particles taking a step and generating a torque in a unit of time; *N*_on_ is the number of particles bound to the rail in the unit of time; *b*_θ_ is the rotational diffusion of the cargo with a normal distribution with zero mean and a standard deviation of *σ*_θ_; *θ*_step_ is the twist angle of the spring driven by the torque *T*_step_ generated by the single particle, assuming that *θ*_step_ is proportional to *T*_step_ (Materials and Methods). Note that the rotational diffusion of the cargo (*b*_θ_) is negligible for the particle formation using Fig. 8 (Materials and Methods).

**Figure 8.**
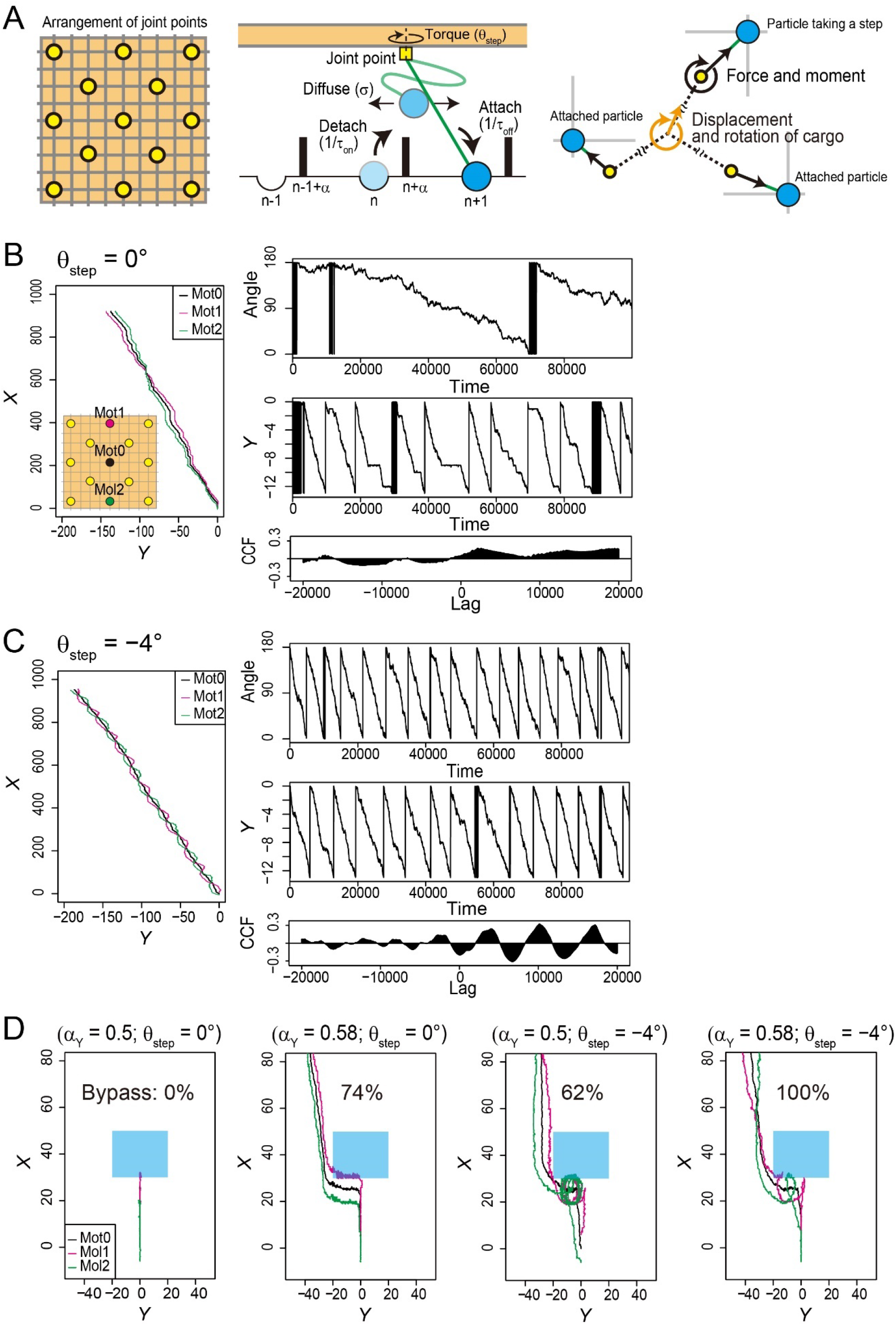
Monte Carlo simulation of the model for kinesin-coated gold nanorods. **(A)** Schematic description of the model. Particles (blue balls) are linked to a rigid body as a cargo (orange square) with a spring (green line), and the formation of their joint points (yellow circle or square) are fixed relative to the cargo axis. Particles repeatedly detach and attach to the 2D lattice rail with rate constants of 1/*τ*_on_ and 1/*τ*_off_, respectively. A detached particle diffuses around the joint points with a standard deviation of *σ*. When the particle passes over the barrier with an asymmetric factor (*α*), at the last time of diffusion, it will attach to the neighboring binding site and generate torque, twisting the spring by *θ*_step_. Translational and rotational motions of the cargo are driven by the balances of force and moment (torque). **(B, C)** Simulation results without and with torque generation (*θ*_step_ = 0° in (B) and *θ*_step_ = −4° in (C)). The parameters are specified in Methods. (left) Typical *X-Y*-trajectories of three representative particles (black, magenta, and green filled circles) represent biased forward, sideward and rotational motions of the cargo. (right) Time trajectories of the angle and *Y*-displacement, and their cross-correlation function (CCF). The axes of the angle and the *Y*-direction are converted to 180°- and 13-lattice periods, respectively. **(D)** Typical examples of simulations of bypassing an obstacle. *α*_Y_ and *θ*_step_ are the sideward asymmetric and torque parameters, respectively (see the main text). The percentages represent the probabilities of success in bypassing the roadblock (blue rectangle) within a calculation time of 55,000 and out of 300 trials.

First, we optimized the parameters for the biased forward and sideward movement of the cargo without torque generation (*θ*_step_ = 0), matching the ~0.8-μm helix pitch of KIF1A and ZEN-4 in the gliding assay, which is the ratio of forward steps to leftward steps (~100/13) (Fig. 8B). We note that the simulation without torque generation often showed unidirectional rotation of the cargo, but the period of rotation was irregular and uncorrelated with the 13-lattice period of the *Y*-displacements modeled as the 13-protofilament microtubule. We then performed the simulation with torque generation. When *θ*_step_ was approximately −3° to −4°, the cargo exhibited periodical clockwise rotation and the period of rotation closely matched the 13-lattice period (Fig. 8C and Fig. S7). Correlation between counterclockwise rotation and sideward movement of the cargo was also reproduced when *θ*_step_ was +4° (Fig. S8). The lateral displacement in a calculation time in the presence of torque (*θ*_step_ = −4°) increased about 1.3-fold, compared with that in the absence of torque (*θ*_step_ = 0°). Taken together, the Monte Carlo simulations are consistent with the analysis by the noise-driven ratchet model showing that unidirectional rotation of a motor-coated cargo enhances the sideward bindings of motor molecules toward the biased direction (Fig. 7).

## Discussion

We developed the GNR motility assay and harnessed it to analyses cargo rotations driven by teams of single-headed KIF1A, dimeric ZEN-4. For these kinesins, ~20% of the kinesin-coated GNRs showed periodic changes in the GNR polarization signal, phase locked to the orbit of the GNR around the microtubule, over multiple orbital periods. For the remaining ~80% of records, we did not observe periodic variation in the GNR polarization signals. The low probability of observation suggests that stable periodic polarization rotations were observed only when the plane of yawing of the kinesin-GNRs about its short axis was nearly parallel to the longitudinal axis of the microtubule. Due to the relatively low aspect ratio of the GNRs we used, the elevation angles of the long axis of the kinesin-GNRs might tend to change, resulting in unstable polarization fluctuations in the majority of the kinesin-GNRs. However, GNRs with higher aspect ratio may inhibit yawing of the kinesin-GNR, since the parallel longitudinal axis alignment of the kinesin-coated GNR and the microtubule is most stable in terms of the number of kinesin molecule interacting with the microtubule.

In the GNR assay experiments, each kinesin-coated GNR exhibited specific directionality of yawing, which is consistent with previous single-molecule measurements for the kinesin-1 dimer(9). Although the detailed mechanism of directional yawing of a kinesin team remains unclear, not only the power stroke in the motor domain but also the formation of kinesin molecules on the GNR and further the configuration of the kinesin-GNR and the microtubule might be involved.

A recent single-molecule study of the kinesin-1 dimer construct revealed that it unidirectionally rotates the stalk domain by ~1° with 8-nm steps at high ATP conditions and in the presence of an external force(9). Therefore, the estimated *θ*_step_ of 4° might be reasonable especially in the absence of an external force. Previous structural studies of kinesin-1 and KIF1A revealed that rearrangement in the motor domain from the pre-stroke state to the post-stroke state generates global rotation of the motor domain(31–33). The rotation of the kinesin motor domain may generate a torque (Supplementary text). The neck-to-tail region is critical for torque transmission from the head domain to the cargo. In the case of ZEN-4, its neck domain contains a non-canonical long sequence; however, high-speed AFM imaging of a ZEN-4 dimer revealed a distinctive globular mass in the neck region(34). Thus, the non-canonical neck-domain of ZEN-4 may be a stiffness component rather than a flexible linker.

### Ideas and speculations

Previous studies proposed that sideward stepping of kinesins and dynein allows them to bypass roadblocks on a microtubule(6–8, 11, 13, 35). We further performed Monte Carlo simulations of bypassing of a roadblock with various multiple-motor motilities, especially changing the two parameters: the off-axis asymmetry, *α*_Y_, and the torque generation, *θ*_step_ (Fig. 8D and Materials and Methods). The simulation results in the case of *α*_Y_ = 0.5 and *θ*_step_ = 0° showed that the motor team failed to bypass it within the calculation time (Fig. 8D, the most left), whilst the off-axis asymmetry (*α*_Y_ = 0.58 and *θ*_step_ = 0°) allow to avoid the roadblock (Fig. 8D, second from the left). Moreover, the simulation results in the case of *α*_Y_ = 0.5 and *θ*_step_ = −4° indicate that the rotational motility also helps the motor team avoid the roadblock (Fig. 8D, second from the right). Interestingly, the motors with off-axis asymmetry (*α*_Y_ = 0.58) and torque generation (*θ*_step_ = −4°) exhibited a ‘spin move’ (i.e., a combination of spinning and sideward motion), and further increased the probability of bypassing the roadblock (Fig. 8D, the most right). The evasion time by the spin move was 2.6-fold faster than that by only sideward stepping. Our theoretical analyses with a noise-driven 2D ratchet model and Monte Carlo simulations showed that unidirectional rotation enhances the biased sideward movement of kinesin-bound cargo and further that the evasion time by the spin move was faster than that by only sideward stepping. Therefore, the spin move of a kinesin team may help sustained transport of a cargo in situations where the microtubule lattice is crowded with microtubule associated proteins, motor proteins, and other lattice-binders. Rotational and translational dynamics of endosomes in cells are not always correlated(36). Distinct changes in the rotational state of endosomes during pausing in translational motion(36) may be related to the spin move of a motor team during sidestepping an obstacle.

In summary, this study reveals that in kinesin teams, the production of torque drives not only orbital motion of the kinesin team around the microtubule axis, but also coupled yaw-axis rotation of the team and its attached cargo. We propose based on quantitative simulations that this combination of properties enables cargos bound to multiple kinesins more efficiently to bypass roadblocks. Our work provides new insights into the structural basis of kinesin motor stepping(33, 37) and into the implications of their translational and rotational degrees of freedom for their cellular function(36).

## Materials and Methods

### Gold nanorod imaging

Fig. 1A shows a schematic drawing of the laser dark-field microscope system used for gold nanorod (GNR) imaging. The laser dark-field optical system was constructed on an inverted microscope (Ti; Nikon, Tokyo, Japan), equipped with variable-angle epi-illumination(21, 22) and a custom-built perforated dichroic mirror (PDM) (Sigma Koki, Tokyo, Japan). The PDM had an elliptical hole (short axis = 6 mm; long axis = 8.5 mm) at its center position, which made a circular window when viewed from the optical axis. The scattered light from the GNRs was transmitted through this central circular window. Illumination was performed using a 635-nm diode laser (MRL-III-635L-10mW; CNI laser, Changchun, China), which was further attenuated by neutral density filters (ND) and converted to circularly polarized light by a quarter-wave plate (WPQ-6328-4M; Sigma Koki, Tokyo, Japan). The collimated and circularly polarized incident beam was focused by a lens (L1 in Fig. 1A, *f* = 220 mm) onto the back focal plane of the objective lens (PlanApo N, 60×, NA 1.42; Olympus, Tokyo, Japan). GNR images were obtained by laser dark-field imaging using highly inclined illumination with nearly total internal reflection angle. The back focal plane was focused outside a camera port of the microscope with an achromatic lens (L3 in Fig. 1A, *f* = 60 mm) to make an equivalent back focal plane for constructing the optics for polarization measurements and 3D tracking.

For polarization measurements, the scattered light from a GNR illuminated by polarized incident beam was further split into two orthogonally polarized components by a polarization beam displacer (BD40; Thorlabs, Tokyo, Japan), which was set in the optical path between the relay optics constructed by two lenses (L3 (*f* = 60 mm) and L4 (*f* = 120 mm) in Fig. 1A)(38). Two separated and polarized light beams were imaged as two spots on a CMOS camera (LDH2500; Digimo, Tokyo, Japan). The orientations of the two orthogonally separated polarization components of the beam displacer were diagonal relative to the transmission axis of the polarization filter. For angular calibration, a linear polarizer was additionally set at the position between L2 and L3 and rotated from 0° to 180° in 10° intervals (Fig. 1B).

For 3D tracking, concave and convex cylindrical lenses (CnC (*f* = −200 mm) and CvC (*f* = 200 nm)) were set in front of the camera(24). The interval between CnC and CvC was 13 mm. Calibration of 3D tracking along the Z-axis was performed by moving the objective in 50-nm intervals (Fig. S1), using a custom-built stage equipped with a pulse motor (SGSP-13ACT; Sigma Koki, Tokyo, Japan) and a controller (QT-CM2; Chuo Precision Industrial, Tokyo, Japan). We analyzed the widths of the GNR spots along the *X*-axis (*S*_X_) and the *Y*-axis (*S*_Y_) obtained by 2D-Gaussian fits. The calibration factor for the z-displacement obtained from the ratio of *S*_Y_ to *S*_X_ was 0.98 μm/ratio.

The temperature of the sample stage was controlled using a custom-built water-circulating temperature control system. Image acquisition in the GNR motility assay was performed at 100 frames/s with the commercial software for the CMOS camera (Digimo, Tokyo, Japan). The data were saved as eight-bit AVI files. The effective pixel size of the detector was 42 nm. Image analysis and further analyses were performed using custom software written in LabVIEW (National instruments, Tokyo, Japan).

### Polarization measurements

For polarization measurements, the scattered light from a GNR illuminated by polarized incident beam was split into two polarized components by the polarization beam displacer as mentioned above. Thus, the intensities of two polarized GNR spots as functions of the GNR orientation (*θ*) are expressed by

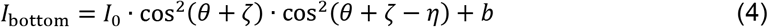

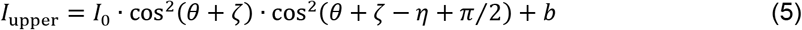

where *I*_0_ is the amplitude; *θ* is the GNR orientation in the sample plane; *ζ* and *η* are phase angles of polarization of the incident beam in the sample plane and the beam displacer, respectively; and *b* is the background intensity. The observed intensity profiles in Fig. 1B were well fitted by these functions with *I*_0_ = 127.5 ± 3.3, *ζ* = 99.3° ± 0.8°, *η* = 45.3° ± 0.7°, and *b* = 19.5 ± 0.9. GNR angles in the sample plane were obtained by fitting an ellipse function to the plot of the degree of polarization (*I*_bottom_ – *I*_upper_)/(*I*_bottom_ + *I*_upper_) as a function of the summed-intensity (*I*_bottom_ + *I*_upper_), as shown in Fig. 1B.

### Kinesin constructs

In all kinesin constructs used in this study, the functional sequence Avi-His, which consists of the Avi-tag (GLNDIFEAQKIEWHE) for biotinylation and the 6×His-tag for histidine-affinity purification, was fused to the C-terminus of truncated kinesin domains. For the monomeric KIF1A construct, the fragment of human KIF1A (1–366 amino acid residues (AAs)) obtained from a human cDNA library was cloned into eGFP-Avi-His/pColdIII plasmids. eGFP stabilized the solubility of the truncated KIF1A sample. In addition, for the surface-gliding assay of the KIF1A construct, we used the construct of human KIF1A (1–366 AAs) cloned into mRuby-Zdk1-His/pColdIII plasmids. For the dimeric ZEN-4 construct, the fragment of C. elegans ZEN-4 (1–555 AAs) obtained from CBD-TEV-ZEN4/pGEX-6p plasmids(34) was cloned into Avi-His/pET32a plasmids. For the monomeric and dimeric kinesin-1 constructs, the fragments of rat kinesin-1 (1–430 AAs for the dimeric constructs(27)) were cloned into Avi-His/pColdIII plasmids. All constructs were verified by DNA sequencing.

### Expression and purification of kinesin

For expression and purification of the kinesin constructs, the proteins were expressed in *Escherichia coli* strain BL21 Star (DE3) cells using 0.4 mM Isopropyl-β-D-thiogalactopyranoside (IPTG) (Sigma) for 9 to12 hours at 20 °C. The collected cells were lysed in lysis buffer (50 mM potassium phosphate; pH 7.4, 350 mM NaCl, 1 mM MgCl_2_, 10 μM ATP, 1 mM DTT, 0.1% Tween20, 10% glycerol, protease inhibitors, 50 mM imidazole) and sonicated in iced water for 15 minutes. The bacterial lysate was centrifuged for 20 minutes at 260,000*g* at 4 °C to remove cell debris and insoluble proteins. The clarified supernatant was purified with the His-tag affinity by chromatography with a HisTrap HP Ni^2+^sepharose column (GE Healthcare). The collected peak fraction was further purified through a HiTrap desalting column (GE Healthcare) to exchange the buffer (20 mM potassium phosphate; pH 7.4, 1 mM MgCl_2_, 1 mM DTT, 10 μM ATP) containing 80 mM NaCl for the KIF1A and the kinesin-1 constructs, or 250 mM NaCl for the ZEN-4 construct.

For the KIF1A construct, we further purified the active proteins based on their affinity for microtubules. KIF1A and microtubules were mixed and incubated for 15 minutes in the presence of 1 mM AMPPNP. The mixture was centrifuged for 20 minutes at 260,000*g* at 25 °C to remove unbound KIF1A molecules. The pellet was resuspended in ATP-containing buffer (20 mM PIPES-KOH; pH 7.4, 5 mM MgCl_2_, 10 mM potassium acetate, 250 mM NaCl, 20 μM paclitaxel, 5 mM ATP) and incubated for 15 minutes at 23 °C for detachment of KIF1A molecules from the microtubules. Microtubules were removed by centrifugation (260,000*g*, 15 minutes, 23 °C).

The purified proteins were flash-frozen and stored in liquid nitrogen. The concentration of proteins was estimated using the Bradford method with the use of BSA as a standard.

### Purification of tubulin

Tubulin was purified from porcine brains through four cycles of temperature-regulated polymerization and depolymerization in a high-molarity PIPES buffer to remove contaminating MAPs(39). The purified tubulin was flash-frozen and stored in liquid nitrogen.

### Preparation of Cy5-microtubules

Tubulin stock solution (~6 mg mL^−1^, 50 μL) and Cy5-labelled tubulin solution (~20 mg mL^−1^, 1 μL, ~50% labelled) were mixed on ice and then diluted twofold with chilled BRB80 buffer (80 mM PIPES-KOH; pH 6.8, 1 mM MgCl_2_, 1 mM EGTA). One microliter each of GTP (100 mM), and MgCl_2_ (100 mM) was added to the diluted tubulin solution. The tubulin solution was incubated at 37 °C for 1 hour for polymerization and stabilized with 20 μM paclitaxel. The Cy5-microtubules were further purified by centrifugation. Ray et al. reported that polymerization in PIPES buffer results in ~60% of 14-protofilament microtubules and ~30% of 13-protofilament microtubules(40).

### Preparation of protein-G-coated beads and streptavidin-coated gold nanorods

For preparation of protein G-coated beads, carboxylate-modified fluorescent microbeads (0.5 μm; Thermo Fisher Scientific, USA) were diluted in the activation buffer (100 mM MES; pH 6.0). EDC and sulfo-NHS were added to the bead solution and incubated for 15 minutes at room temperature. The beads were then washed with 100 mM HEPES buffer at pH 8.0. Protein-G was then added to the bead solution and reacted for 2 hours at room temperature. Any unreacted amine reactive groups were quenched by adding excess glycine. Excess proteins were removed by centrifugation (21,000*g*, 15 minutes, 23 °C). The beads were resuspended in BRB80 buffer.

For preparation of streptavidin-coated GNRs, biotin-labelled GNRs (5 nM) (C12-40-600-TB; Nanopartz, CO, USA) and streptavidin (5 mg mL^−1^) were mixed. After 15 minutes’ incubation, excess streptavidin was removed by centrifugation (4,000*g*, 3 minutes, 23 °C) and then the pellet was resuspended in Milli-Q water and sonicated for 2 s. The procedure for removing excess streptavidin was repeated twice.

### GNR motility assays

Streptavidin-coated GNR (2 μL, 5 nM) and kinesin sample (2 μL, 1-2 μM) were mixed and incubated for at least 15 minutes at room temperature. For the assay of KIF1A, 2 μL of 0.2% Tween20 was further added to the mixture of kinesin and GNR. For the assay for ZEN-4, 0.5 μL of 5 M NaCl and 2 μL of 0.2% Tween20 were further added to the mixture of kinesin and GNR.

A flow chamber (sample volume of about 5 μL) was prepared using two coverslips (24 mm × 32 mm and 18 mm × 18 mm) stuck by double-sided tape. Protein-G-coated bead solution was nonspecifically absorbed to the coverslip. After 3 minutes’ incubation, the flow chamber was washed with 20 μL of 1 mg mL^−1^ BSA and further incubated for 3 minutes. The flow chamber was washed with 20 μL of BRB80. Then, 10 μL of 20 μg mL^−1^ anti-β-tubulin antibody solution (sc-58884; Santa Cruz, USA) was flowed to the flow chamber. After 3 minutes’ incubation, the flow chamber was washed with 20 μL of BRB80. Then, 10 μL of Cy5-lablled microtubule solution (~2 μM tubulin concentration) was flowed into the flow chamber such that the microtubules bound to the beads (Fig. 1C). After 3 minutes’ incubation, the flow chamber was washed with 20 *μ*L of 1 mg mL^−1^ casein. After 3 minutes’ incubation, the flow chamber was washed with 20 μL of the motility buffer (20 mM PIPES-KOH; pH 7.4, 4 mM MgCl_2_ and 10 mM potassium acetate) containing the kinesin-coated GNRs (~100 pM), ATP (2 mM), the ATP regeneration system (0.8 mg mL^−1^ creatine kinase, 10 mM creatine phosphate), the oxygen scavenger system (3 mg mL^−1^ glucose, 50 U mL^−1^ glucose oxidase, 1200 U mL^−1^ catalase), and 1 mg mL^−1^ casein. The GNR assays were performed at 27 °C. Image acquisition was performed at 100 frames s^−1^.

### Surface-gliding assays for KIF1A with QD-coated microtubules

Qdot525 (QD)-coated and Cy5-labelled microtubules were prepared as previously described(18). The surface-gliding assays were performed in flow chambers assembled from two coverslips attached using double-sided tapes. Before the assay, kinesin stock solution was diluted to 1 μM with motility buffer. Five microliters of a 5 mg mL^−1^ solution of protein G (Sigma-Aldrich, Tokyo, Japan) was added to the flow chamber (about 5 μL). After 5 minutes’ incubation, the flow chamber was washed with 20 μL of BRB80 buffer, followed by another 5 minutes of incubation with 5 μL of 0.05 mg mL^−1^ anti-His-tag antibody. The flow chamber was washed with 20 μL of BRB80 buffer, and then 5 μL of 0.5 mg mL^−1^ casein solution was added. After 5 minutes’ incubation, the flow chamber was washed with 20 μL of BRB80 buffer. Then, 10 μL of KIF1A solution (1 μM) was added to the flow chamber. The flow chamber was then washed with 20 μL of BRB80 after 5 minutes’ incubation, and 20 μL of QD-coated microtubule solution (0.05-0.1 μM tubulin) was added to the flow chamber. Finally, the flow chamber was washed with 20 μL of BRB80 buffer after 5 minutes’ incubation, and 20 μL of motility buffer containing 3 mM ATP, 20 μM paclitaxel, 0.5% β-mercaptoethanol, 0.25 mg mL^−1^ casein, ATP regeneration and oxygen scavenger systems were added. Fluorescent images of the QD- and Cy5-labelled microtubules were captured using TIRF microscopy at 25 °C. Corkscrewing pitches in the surface-gliding assay were determined by measuring the longitudinal displacement of the QDs in each period of oscillation of the lateral displacement along the microtubule axis.

### Surface-gliding assays for ZEN4-GNR

Preparations of Cy5-labelled microtubules and ZEN4-GNR were prepared as described above. Surface-gliding assays were performed in flow chambers assembled from two coverslips attached using double-sided tape. The ZEN4-GNR solution was diluted to ~50 pM with motility buffer and added to the flow chamber (about 5 μL). After 5 minutes’ incubation, the flow chamber was washed with 20 μL of BRB80 buffer, followed by 5 minutes’ incubation with 10 μL of 0.5 mg mL^−1^ casein solution. The flow chamber was then washed with 20 μL of BRB80 buffer, and 20 μL of Cy5-labelled microtubule solution (2 mM ATP, Cy5-labelled microtubules, 0.25 mg mL^−1^ casein, and ATP regeneration and oxygen scavenger systems in motility buffer) were added. The assays were performed at 27 °C. Image acquisition of the Cy5-labelled microtubules was performed at 10 frames s^−1^.

### Quantification and statistical analysis

Polarization and position data of the observed kinesin-coated GNRs were obtained by image analysis with software written in LabVIEW (National instruments, Tokyo, Japan). Velocities of the kinesin-coated GNRs were determined by fitting the trace of the longitudinal displacement along the microtubule axis (*X*-direction) with a linear function. Rotation and helix pitches in the GNR assays were determined by measuring the *X*-displacements in the periods of oscillations of the angle- and *Y*-trajectories, respectively. Statistical hypothesis testing was performed using *R* statistical software.

### Correction of the measured pitch

The measured pitch (*P*_Measured_) comprised two helical elements: the helix pitch exerted by motors (*P*_Motor_) without supertwist of the microtubule, and the supertwist of the microtubule depending on the number of the protofilaments (*P*_MT_). *P*_Measured_ is given by

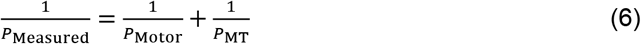

The PMT of 12-, 13-, and 14-protofilament microtubules is estimated to be about −3.4, −24.8, and 6.8 μm, respectively (we assign a positive value to the pitch in the left-handed helix)(40). When the absolute value of *P*_Measured_ is much smaller than the absolute value of *P*_MT_ (< ~1 μm), *P*_Motor_ is not appreciably changed by the supertwist (Fig. S4).

### Mathematical model of helical and rotational motion of a kinesin-coated GNR on a microtubule

The GNR motility assay revealed that KIF1A-, ZEN4-, and sK1-GNRs unidirectionally rotate about the GNR’s short axis, moving along the left-handed helix track. The helix track of the center position of a GNR is given by

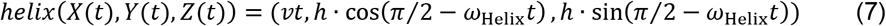

where *v* is the translational velocity along the *X*-axis, *ω*_Helix_ is the angular velocity in the *Y-Z*-plane, and *h* is the radius of the helix. Assuming that the long axis of the GNR is parallel to the *X-Y*-plane under the initial conditions, the initial position matrix *p*_0_ consisting of the tip, center, and end positions of the GNR is given by

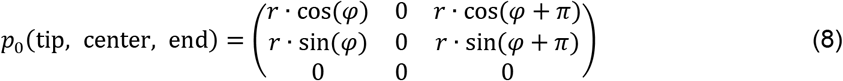

where *r* is the radius along the long axis of the GNR and *φ* is its phase angle. Because a kinesin-coated GNR unidirectionally rotates about its short axis and further around the microtubule, the resultant rotation comprises yawing and rolling. We assume that a kinesin-coated GNR would not rotate about the pitch axis. Matrices of yawing and rolling are given by

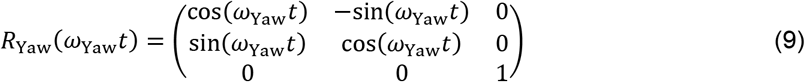

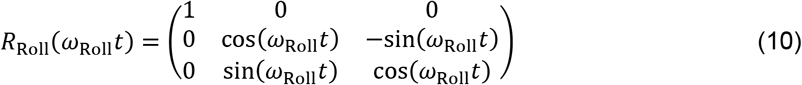

where *ω*_Yaw_ and *ω*_Roll_ are the angular velocities of yawing and rolling, respectively. Taken together, the helical and rotational trajectory of a motor-coated GNR is written as

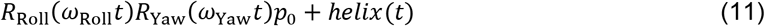

By calculating *R*_Roll_*R*_Yaw_*p*_0_, the time trajectory of the GNR orientation in the *X-Y*-plane is obtained (Equation (1)). Our analyses indicate that kinesin-coated GNRs rotate 180° in one period of the helix. This is the case of *ω*_Helix_ = *ω*_Roll_ = ±2*ω*_Yaw_. The calculated helical and rotational trajectories of a kinesin-coated GNR and the time trajectories of the GNR orientation with different phase angles are shown in Fig. 4, Fig. S6, and Movies S2, S4, and S5.

### Evaluation of particle flux in the noise-driven ratchet model

The previous study by Astumian and Bier(30) clarified the particle flux in the two-state, noise-driven ratchet model. Following their discussions, we evaluated the particle flux for different start positions, because sideward translations (±*d*) of motors takes place owing to the rotation of their cargo (see Fig. 7A). In the two-state ratchet model, when the barrier moves to the down state, the particles start to diffuse. The probability density function of a diffusing particle at the start position of +*d* in the *X*-direction is given by

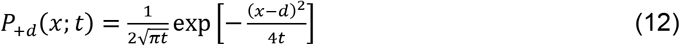

After 1/*γ* units of time, the probability that the particle is located to the right of *α* is given by

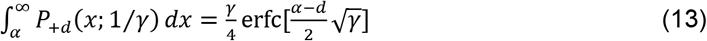

and the probability that it on the left of −1+*α* is given by

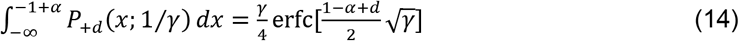

The particle flux, *J_+d_* (Equation (3)) is given by equation (13) and equation (14). Similarly, *J_−d_* (Equation (3)) and *J*_0_ (Equation (2)) are also obtained.

Following the probability density function of a diffusing particle, the standard deviation of diffusion is given by 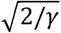. When *γ* is 160 s^−1^, 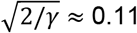, which is equal to the value of *σ* in the Monte Carlo simulations (see below).

### Monte Carlo simulations

We performed Monte Carlo simulations to model the observed rotational and helical motion. In the simulation, each particle is linked to a rigid body as a cargo with a spring (spring constant, *k*) and the joint points linking the particles and the cargo are fixed. The particles stochastically attach and dissociate from the rail (on-time constant, *τ*_on_; off-time constant, *τ*_off_). A detached particle diffuses around the joint point (standard deviation of diffusion, *σ*) in the *X-Y*-plane. When the diffusing particle overrides the asymmetric barrier (asymmetric factors, *α*_X_ and *α*_Y_) at the last time of diffusion, the particle takes a step in the *X-Y*-direction. The force- and moment-balance equations are given by 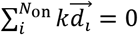 and 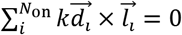, where *k* is the spring constant; 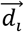 is an extension vector of the spring of the *i*-th particle that is bound to the rail; 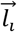 is the position vector of the *i*-th joint relative to the center of the joint points connecting the particles that are bound to the rail; and *N*_on_ is the number of the particles bound to the rail. In the simulation, 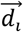 is the displacement between the *i*-th particle and the *i*-th joint position. Furthermore, torque rotating the cargo is exerted by the particle when it takes a step. A twist angle (Δ*θ*) of the cargo by the torque generated by the particle taking a step is given by Δ*θ* = (*N*_step_*θ*_step_ + *b*_θ_)/*N*_on_, and *θ*_step_ = *T*_step_*L/KG*, where *N*_step_ is the number of particles taking steps; *N*_on_ is the number of particles bound to the rail; *θ*_step_ is the twist angle of the spring; *b*_θ_ is the rotational diffusion of the cargo as a normal distribution with zero mean and a standard deviation of *σ*_θ_; *T*_step_ is the torque force generated by the single particle; *L* is the length of the spring; *G* is the modulus of rigidity of the spring; and *K* is the polar moment of inertia of area of the spring. We thus assume that the twist angle *θ*_step_ is proportional to the torque force *T*_step_. The value of *θ*_step_ as the torque force was given in the simulation. After calculating the force and moment balance equations, the cargo additionally rotates Δ*θ* degrees about the center of the joint points connecting the particles that are bound to the rail.

A roadblock in the simulation allowed the off-state particles to diffuse above it, whereas the on-state particles could not attach to the roadblock except at its edges. We used the roadblock with a width of 40 lattices and a length of 30 lattices.

The simulation was developed in LabVIEW (National Instruments). LabVIEW source code and executable software for the Monte Carlo simulations used in this study are available from GitHub (https://github.com/MitsuSGW/MonteCarlo-MotorTeam-Sim.git). The values for the parameters used in the simulation were: *τ*_on_ = 40 units of time; *τ*_off_ = 10 units of time; *α*_X_ = 0.12 units of lattice size; *α*_Y_ = 0.58 units of lattice size; *σ* = 0.11 units of lattice size; *θ*_step_ = 4°; *b*_θ_ = 0. The value of *σ* was determined by the evaluation of particle flux in the noise-driven ratchet model mentioned in the previous subsection. The rotational diffusion of the cargo (*b*_θ_) is negligible in the case of the particle formation used in Fig. 8 (Fig. S9).

### Statistical analysis

Statistical analysis of two groups was performed with Welch’s two-tailed *t* test. Statistical analysis to compare the observed frequencies of two categories was performed with two-sided binomial test of the null hypothesis that the probability of success in a Bernoulli experiment is 0.5. *P* < 0.05 was considered as statistically significant. Values are represented as means ± SD as indicated in the figures.

## Author Contributions

M.S. and J.Y. designed research; M.S. performed experiments, analyzed data, and performed simulations; Y.M., and M.Y. contributed new reagents; M.S., R.A.C. and J.Y. wrote the paper.

## Acknowledgments

CBD-TEV-ZEN4/pGEX-6p plasmid was a gift from Prof. Masanori Mishima. We thank Prof. TOYOSHIMA Y. Yoko and SAITO Kei for critical discussion; Adam Brotchie, PhD, from Edanz Group (www.edanzediting.com/ac) for editing a draft of this manuscript. This work is supported by Ministry of Education, Culture, Sports, Science and Technology Japan (MEXT) grants KAKENHI JP19K06593 and JP19H03190 (to M.S.), JP20K06635 (to J.Y.): MEXT grants KAKENHI Grant-in-Aid for Scientific Research on Innovative Areas JP21H00386 and JP21H05868 (to J.Y.): Research Foundation for Opto-Science and Technology Japan (to M.S).

## Supplementary Information for

### Supplementary Information Text

#### Torque-generation property built into the kinesin motor domain

The rotational work was estimated to be ~30 pN nm in one ATPase cycle of single-molecule kinesin-1(1); thus, the torque *T*_step_ for a *θ*_step_ of 4° can be estimated to be 430 pN nm (≈ 30 pN nm/0.07 rad). In solid mechanics, the twist angle of an object is given by *θ*_step_ = *T*_step_*L*/(*KG*), where *T*_step_ is the torque, *L* is the length of the spring, *G* is the modulus of rigidity of the spring, and *K* is the polar moment of inertia of the object. Because kinesin molecules were bound to streptavidin molecules attached to a gold nanorod, the motor molecule could be roughly approximated as a cylinder with a diameter (*d*) of ~5 nm and length (*L*) of ~12 nm. The polar moment of inertia of a cylinder is given by *K* = (π*d*^4^)/32. Finally, we obtained the modulus of rigidity of the spring *G* (= *T*_step_*L*/(*Kθ*_step_)) of ~1.2 GPa. The Young’s modulus of the spring is thus estimated to be ~3 GPa at a Poisson ratio of 0.25, which are comparable to the values for actin, tubulin, and coiled-coil (~2 GPa) (2). Therefore, the torque may be generated by conformational changes in the kinesin motor domain.

**Figure S1.**
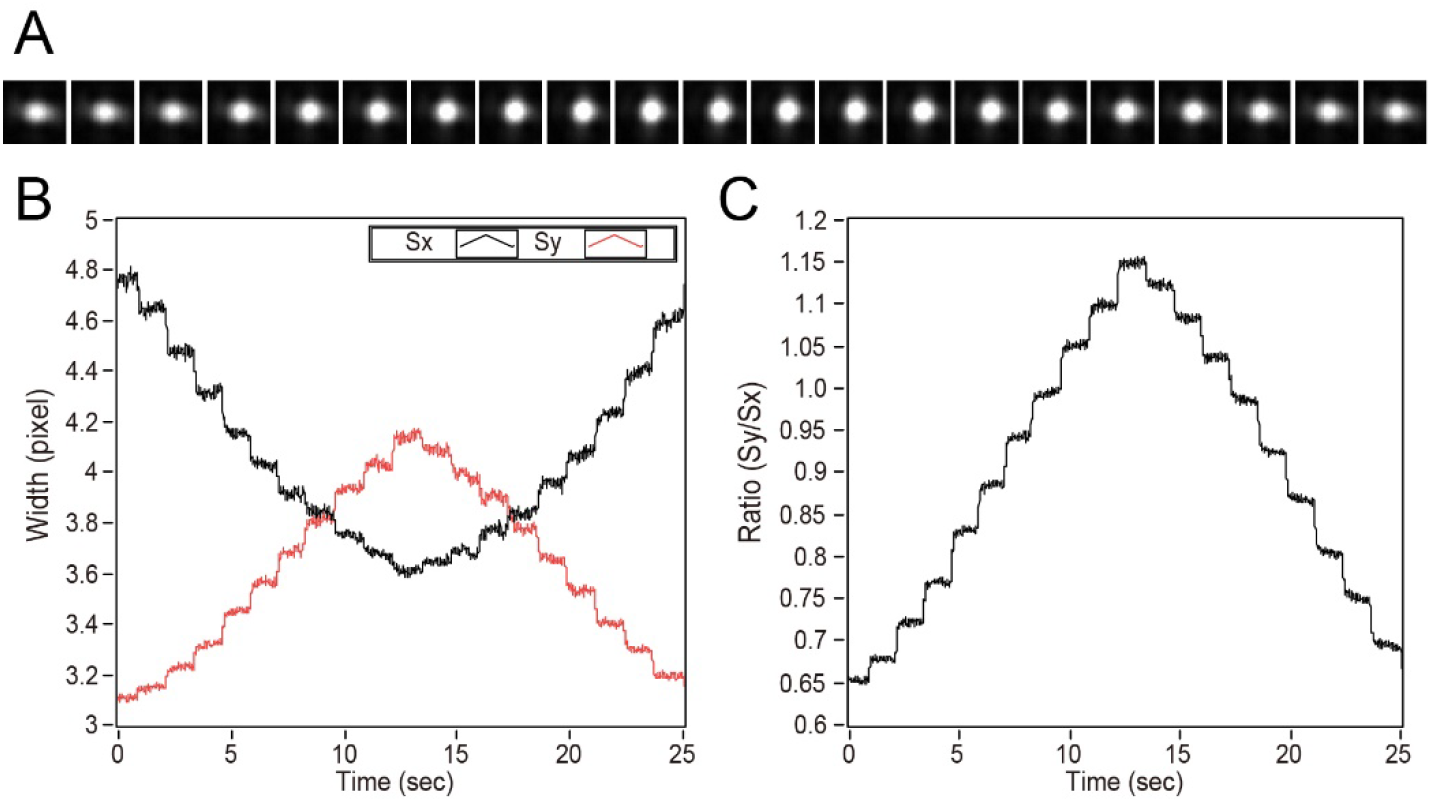
Calibration of GNR displacements along the *Z*-axis. (**A**) Scattered images of a single GNR bound to the coverslip, taking 50-nm steps along the Z-axis. (**B**) Time trajectories of the widths of the GNR spot images in (A) along the *X*-axis (*S*_X_, black line) and *Y*-axis (*S*_Y_, red line). The widths were the standard deviations obtained by 2D Gaussian fits. (**C**) Time trajectory of the ratio of *S*_Y_ to *S*_X_, which represents the *Z*-displacement of the GNR.

**Figure S2.**
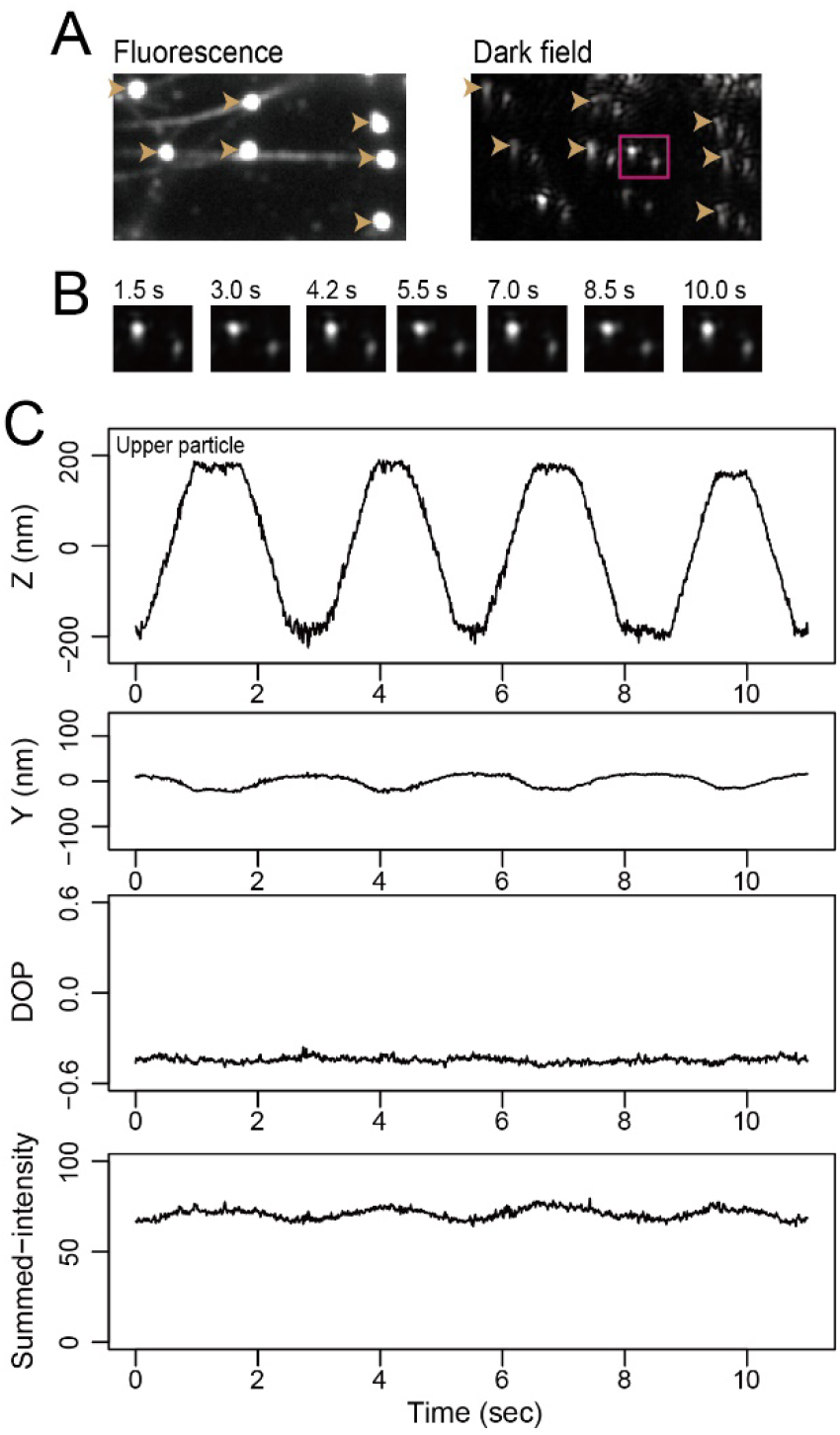
Stability of gold-nanorod (GNR) polarization against the *Z*-displacement. (**A**) Fluorescent image of antibody-coated microbeads (arrowheads) and microtubules, and dark-field image of the microbeads (arrowheads) and the GNRs coated with kinesin-1 (Kin1-GNR) (magenta region of interest). Microtubules were suspended on the microbeads via antigen-antibody system for the β tubulin. The single Kin1-coated GNR was bound on the suspended microtubule in the presence of 1 mM AMPPNP. (**B**) Montage of one pair of the scattered images of the single Kin1-coated GNR shown in the magenta region of interest in (A). The objective lens repeatedly moved ±400-nm along the *Z*-axis. (**C**) Time trajectories of the *Z*- and *Y*-displacements, the degree of polarization (DOP), and the summed-intensity of the single Kin1-coated GNR shown in (B). The *Z*-displacement trajectory was obtained from the upper-left spot in (B).

**Figure S3.**
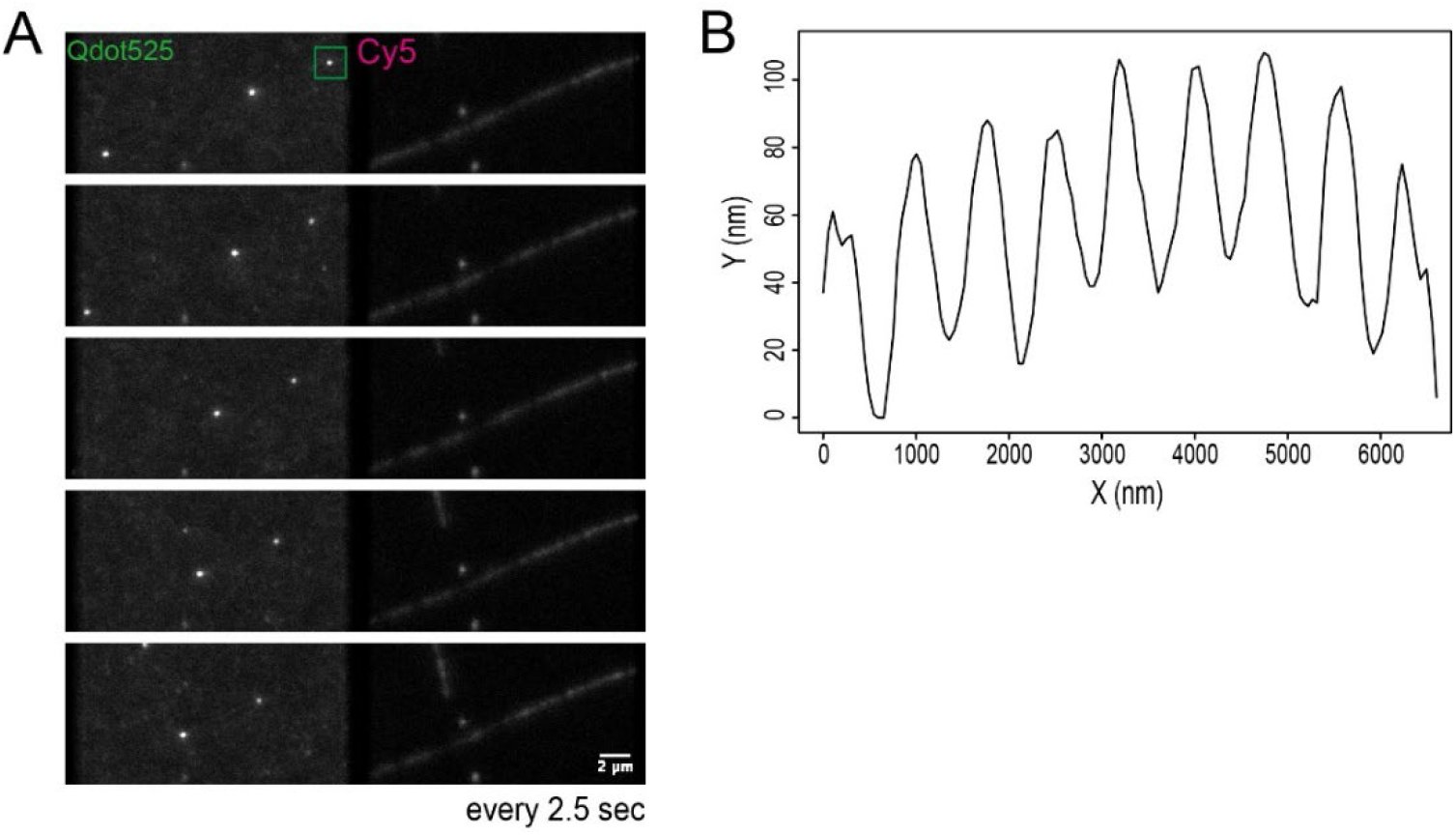
Surface-gliding assay of KIF1A. (**A**) Fluorescence images of Qdot525s and a Cy5-labelled microtubule in the surface-gliding assay of KIF1A. (**B**) Trajectory of the Qdot525 spot in the region of interest in (A), which exhibits a stable sinewave, suggesting the repetitive corkscrewing motions of the microtubule. A corkscrewing pitch of the microtubule was determined by measuring the *X*-displacement in each period of oscillation along the *Y*-axis. The translational velocity was 0.49 ± 0.02 μm/s (mean ± SD, *n* = 5 microtubules) and the corkscrewing pitch was 0.89 ± 0.21 μm (mean ± SD, *n* = 35 revolutions).

**Figure S4.**
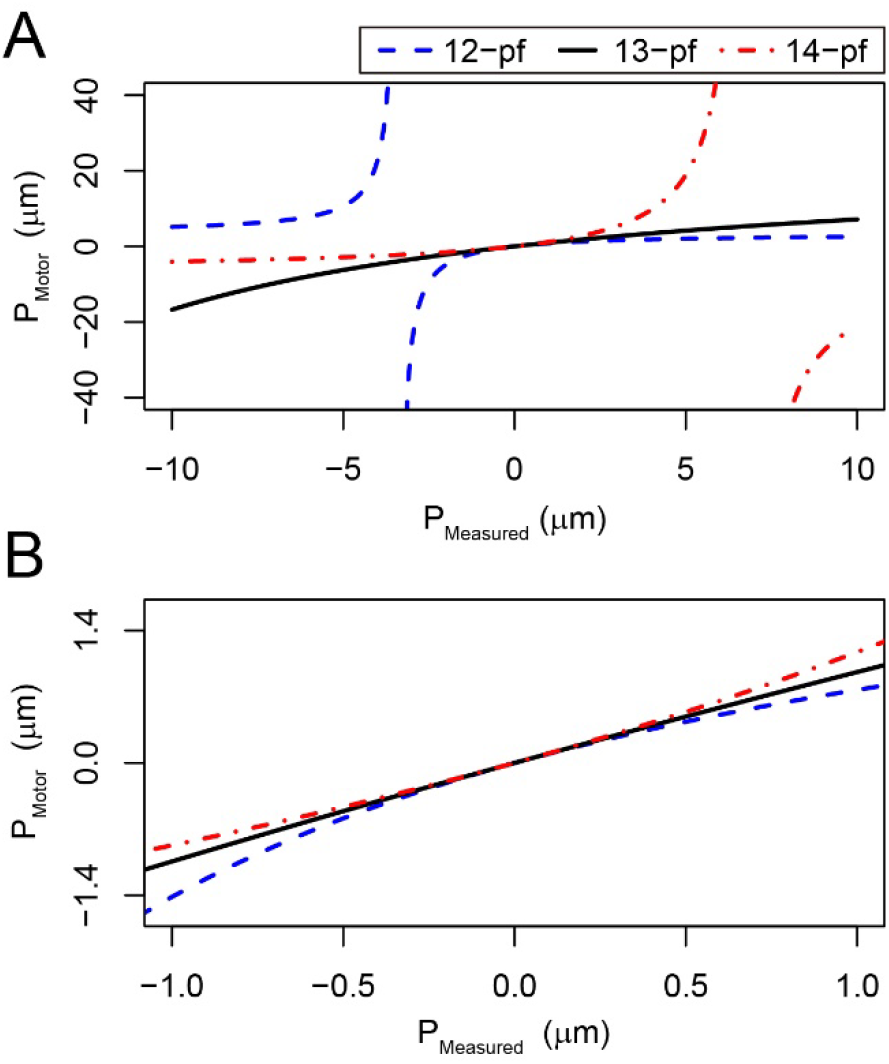
Relationship between measured pitches, pitches of motors and supertwist of microtubules. (**A, B**) Pitches of motors (*P*_Motor_) as a function of measured pitch (*P*_Measured_) and supertwists (*P*_MT_) of 12-, 13-, and 14-protofilament microtubules (12-pf, 13-pf, and 14-pf, respectively). *P*_Motor_ is given by *P*_Motor_^−1^ = *P*_Measured_^−1^– *P*_MT_^−1^. The dashed blue line, the black line, and the doted-and-dashed red line represent the calculation for 12-pf (*P*_MT_ = −3.4 μm), 13-pf (*P*_MT_ = −24.8 μm), and 14-pf (*P*_MT_ = 6.8 μm), respectively.

**Figure S5.**
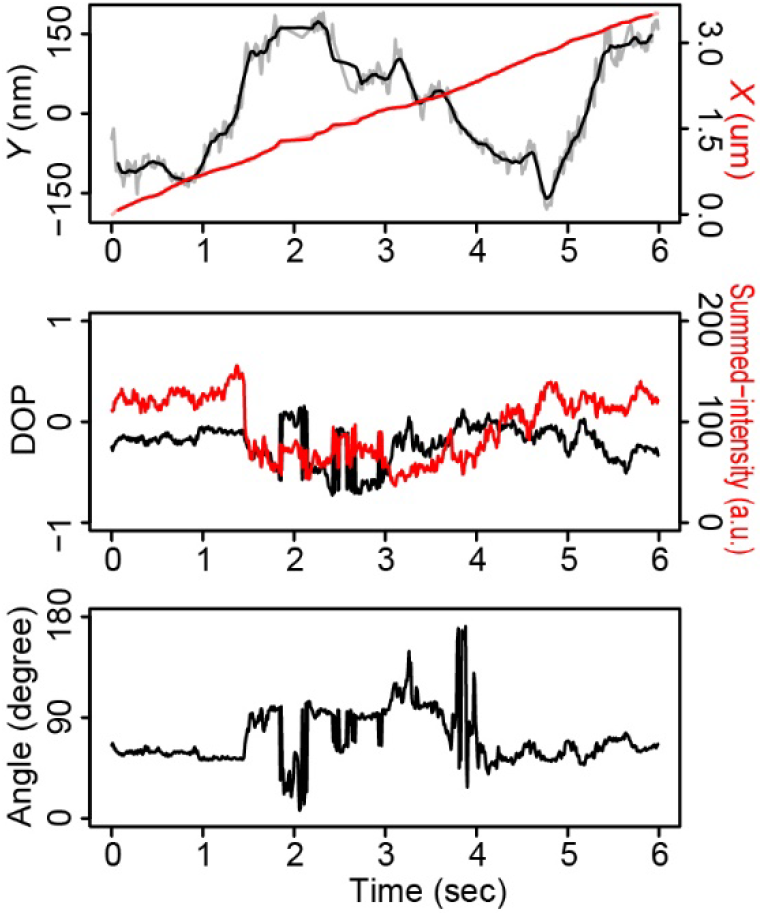
Typical time trajectories of kinesin-1 coated GNR. Time trajectories of the *X*- and *Y*-displacements (upper), the DOP and summed-intensity of the two spots (middle), and the estimated angle (bottom) of the kinesin-1 coated GNR.

**Figure S6.**
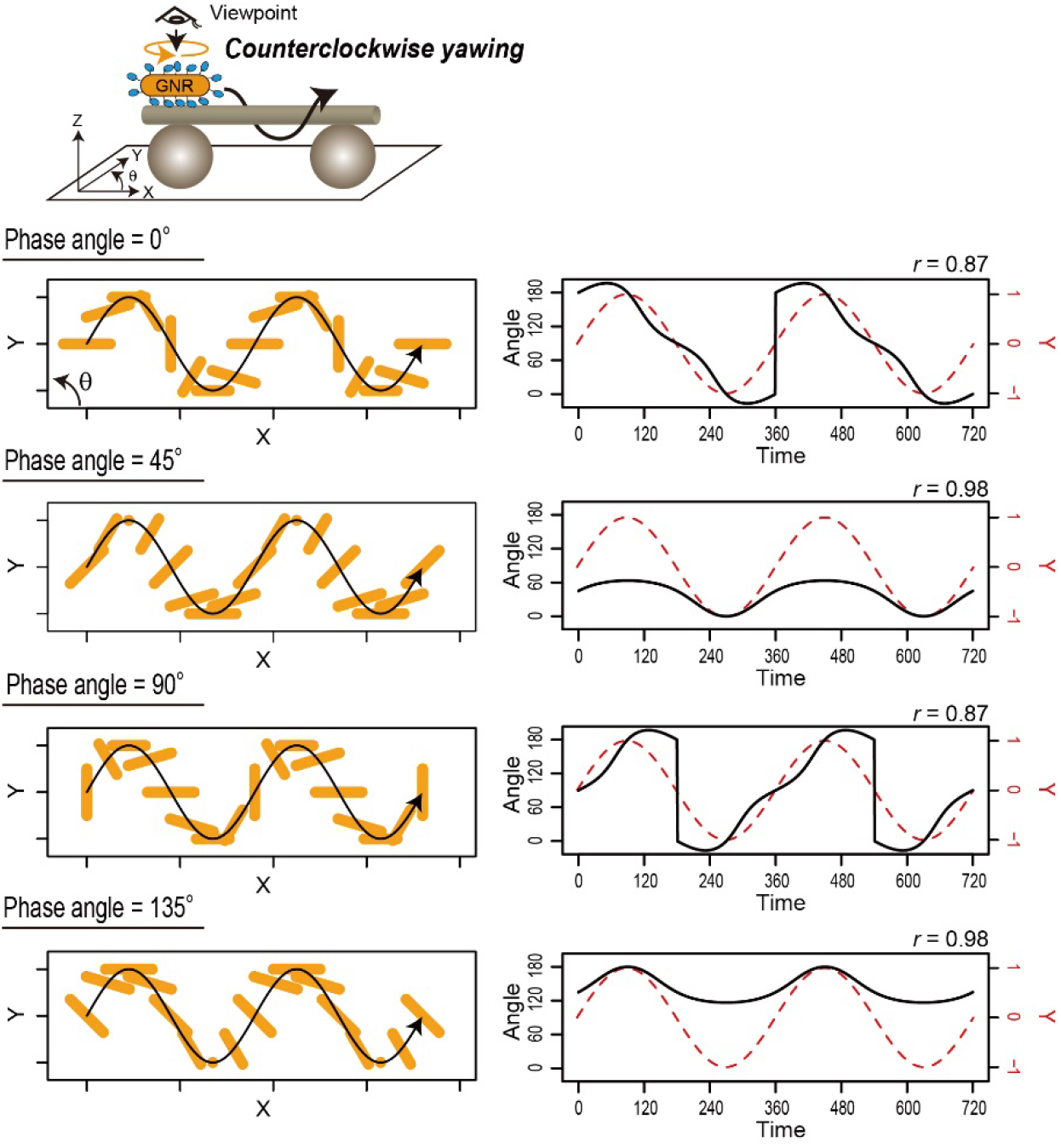
Calculated trajectories of a motor-coated GNR in the case of counterclockwise yawing by the proposed model. Calculation of time trajectories of the angle and the *X-Y*-trajectories of a motor-coated GNR using the proposed model with different phase angles (0°, 45°, 90°, and 135°), in which the kinesin-coated GNR rotates 180° counterclockwise about its yaw axis in one period of helical trajectory. See also Movie S5.

**Figure S7.**
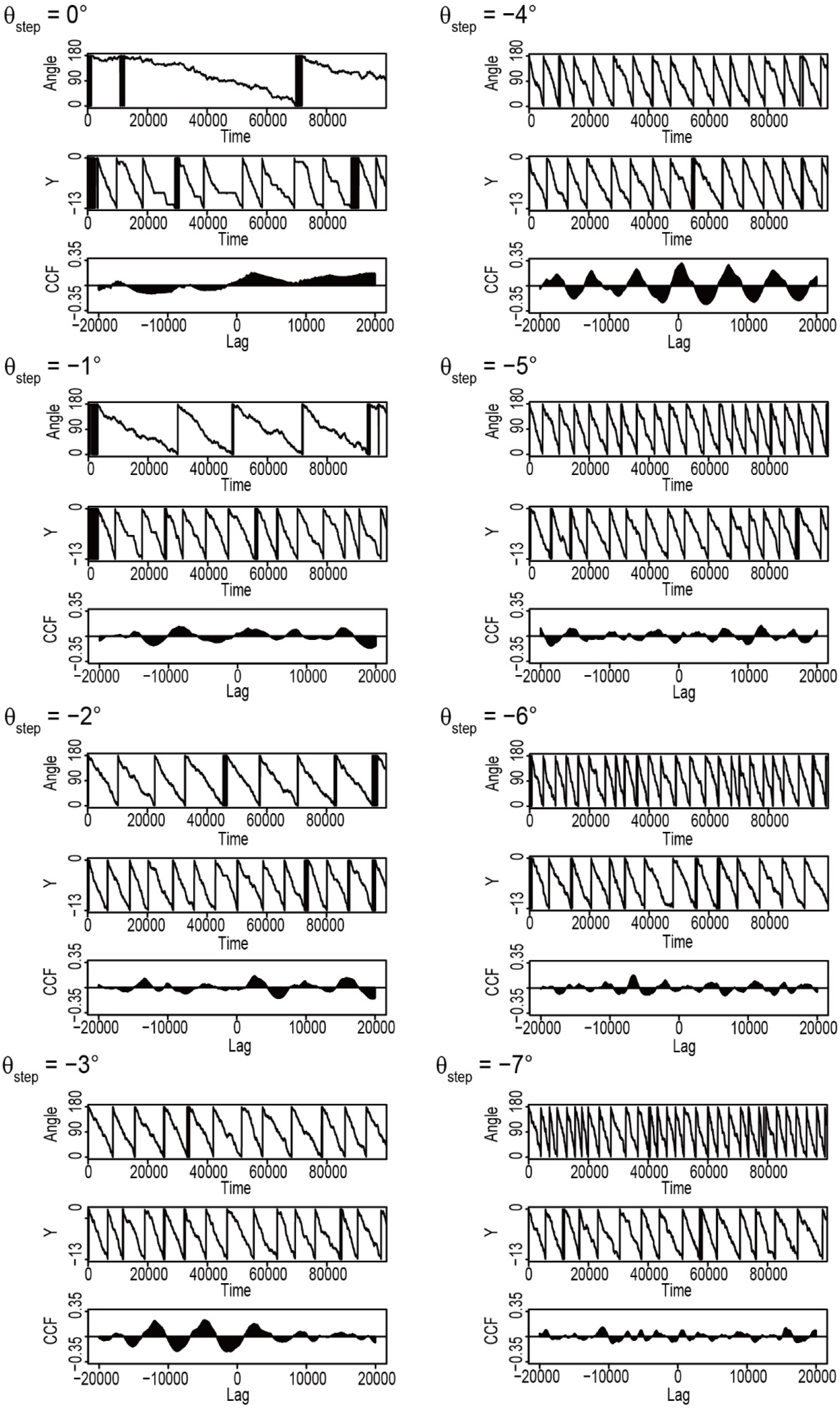
Typical results of the Monte Carlo simulations. Time trajectories of angle of the cargo, the 13-period displacement along the *Y*-axis, and the cross-correlation functions (CCFs) between them, in the range of *θ*_step_ from 0° to −7°.

**Figure S8.**
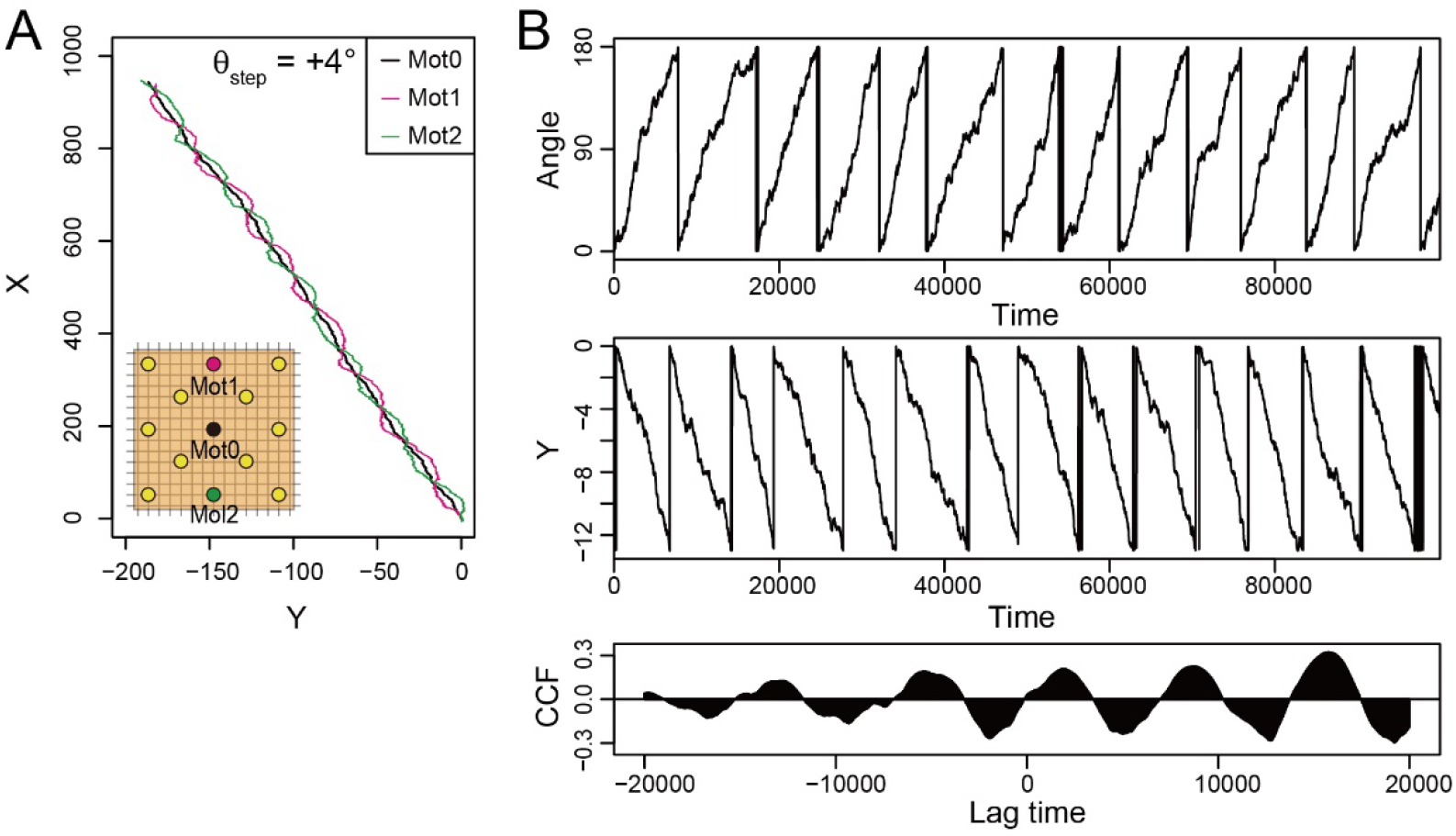
Simulation results with torque generation (*θ*_step_ = +4°). (**A**) Typical *X-Y*-trajectories of three particles (magenta, black, and green filled circles). (**B**) Time trajectories of the angle and the *Y*-displacement of the cargo and their cross-correlation function (CCF), corresponding to (A). The axes of the *Y*-direction are converted into a 13-lattice period.

**Figure S9.**
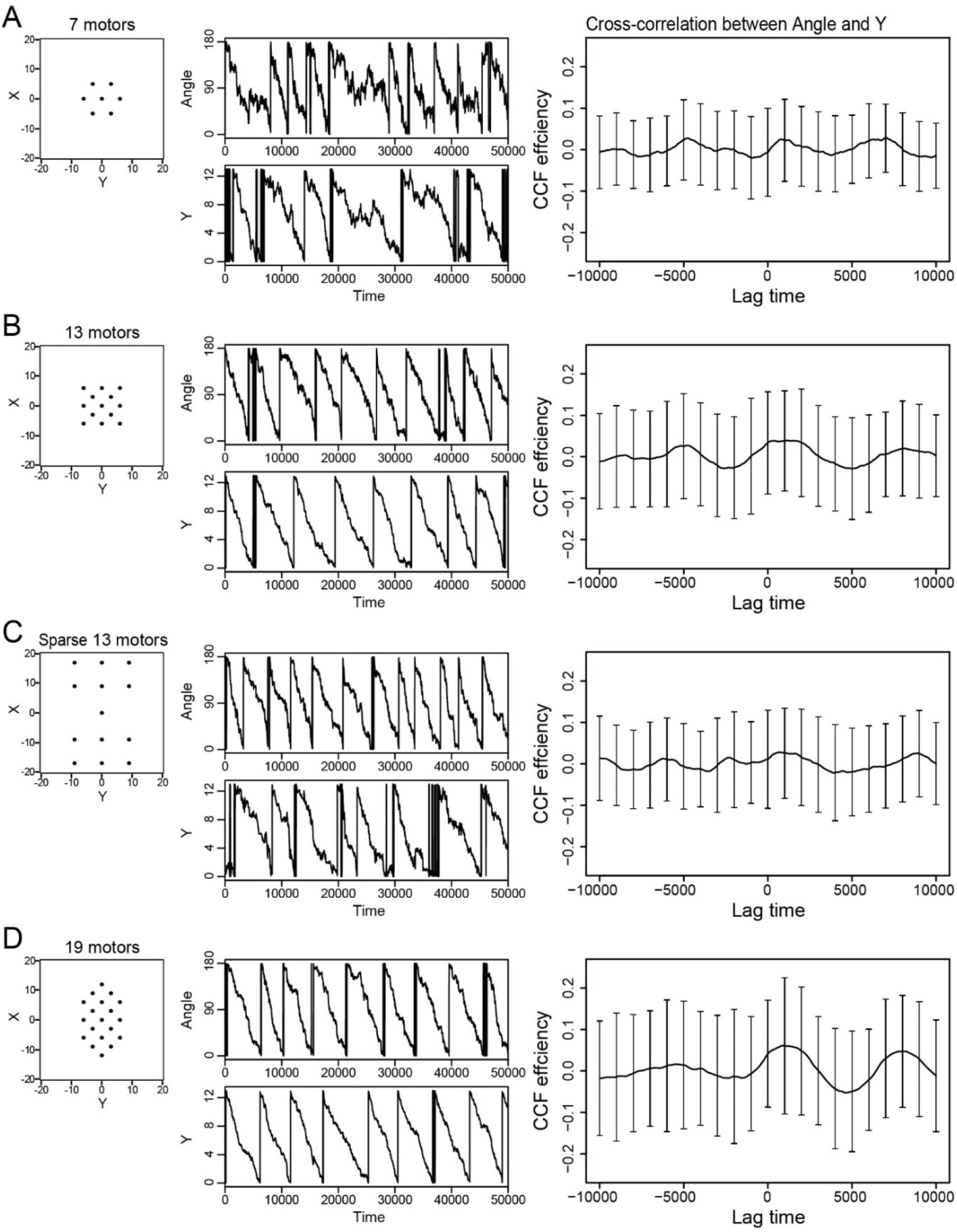
Simulation results with torque generation (*θ*_step_ = −4°) and additional noise (*σ*_θ_ = 8°) in various formations of particles on a cargo. (**A-D**) (left) formations of particles bound on a cargo at initial time. Black dots represent the positions of the particles. (middle) Typical time trajectories of the angle and the *Y*-displacement of the cargo. The axis of the *Y*-direction is converted into a 13-lattice period. (right) Averaged cross-correlation efficiency (CCF) between the angle and the *Y*-displacement (*n* = 100 simulations). Error bars represent standard deviation.

**Movie S1 (separate file).** Corresponds to Fig. 2A.

**Movie S2 (separate file).** Corresponds to Fig. 4A.

**Movie S3 (separate file).** Animation movie of the proposed movement of a kinesin-coated gold nanorod on the microtubule.

**Movie S4 (separate file).** Corresponds to Fig. 4B.

**Movie S5 (separate file).** Corresponds to Fig. S6.

**Movie S6 (separate file).** Corresponds to Fig. 6.

## References

1. R. D. Vale, The molecular motor toolbox for intracellular transport. Cell 112, 467–480 (2003).

2. N. Hirokawa, Y. Noda, Y. Tanaka, S. Niwa, Kinesin superfamily motor proteins and intracellular transport. Nat. Rev. Mol. Cell Biol. 10, 682–696 (2009).

3. K. J. Verhey, J. W. Hammond, Traffic control: Regulation of kinesin motors. Nat. Rev. Mol. Cell Biol. 10, 765–777 (2009).

4. J. Yajima, K. Mizutani, T. Nishizaka, A torque component present in mitotic kinesin Eg5 revealed by three-dimensional tracking. Nat. Struct. Mol. Biol. 15, 1119–21 (2008).

5. B. Nitzsche, F. Ruhnow, S. Diez, Quantum-dot-assisted characterization of microtubule rotations during cargo transport. Nat. Nanotechnol. 3, 552–556 (2008).

6. M. Brunnbauer, et al., Torque Generation of Kinesin Motors Is Governed by the Stability of the Neck Domain. Mol. Cell 46, 147–158 (2012).

7. M. Bugiel, E. Schäffer, Three-Dimensional Optical Tweezers Tracking Resolves Random Sideward Steps of the Kinesin-8 Kip3. Biophys. J. 115, 1993–2002 (2018).

8. A. Mitra, F. Ruhnow, S. Girardo, S. Diez, Directionally biased sidestepping of Kip3/kinesin-8 is regulated by ATP waiting time and motor–microtubule interaction strength. Proc. Natl. Acad. Sci. 115, E7950–E7959 (2018).

9. A. Ramaiya, B. Roy, M. Bugiel, E. Schäffer, Kinesin rotates unidirectionally and generates torque while walking on microtubules. Proc. Natl. Acad. Sci. 114, 10894–10899 (2017).

10. A. Yildiz, M. Tomishige, A. Gennerich, R. D. Vale, Intramolecular strain coordinates kinesin stepping behavior along microtubules. Cell 134, 1030–41 (2008).

11. Y. Maruyama, et al., CYK4 relaxes the bias in the off-axis motion by MKLP1 kinesin-6. Commun. Biol. 4 (2021).

12. X. Pan, S. Acar, J. M. Scholey, Torque generation by one of the motor subunits of heterotrimeric kinesin-2. Biochem. Biophys. Res. Commun. 401, 53–57 (2010).

13. V. Bormuth, et al., The highly processive kinesin-8, Kip3, switches microtubule protofilaments with a bias toward the left. Biophys. J. 103, L4–L6 (2012).

14. R. A. Walker, E. D. Salmon, S. A. Endow, The Drosophila claret segregation protein is a minus-end directed motor molecule. Nature 347, 780–782 (1990).

15. B. Nitzsche, et al., Working stroke of the kinesin-14, ncd, comprises two substeps of different direction. Proc. Natl. Acad. Sci. 113, E6582–E6589 (2016).

16. A. Mitra, et al., Kinesin-14 motors drive a right-handed helical motion of antiparallel microtubules around each other. Nat. Commun. 11, 1–11 (2020).

17. J. Yajima, R. A. Cross, A Torque Component in the Kinesin-1 Power Stroke. Nat. Chem. Biol. 1, 338–341 (2005).

18. M. Yamagishi, S. Fujimura, M. Sugawa, T. Nishizaka, J. Yajima, N-terminal β-strand of single-headed kinesin-1 can modulate the off-axis force-generation and resultant rotation pitch. Cytoskeleton 77, 351–361 (2020).

19. A. Mitra, et al., A Brownian ratchet model explains the biased sidestepping of single-headed kinesin-3 KIF1A. Biophys. J. 8, 1–9 (2019).

20. M. Yamagishi, Y. Maruyama, M. Sugawa, J. Yajima, Characterization of the motility of monomeric kinesin-5/Cin8. Biochem. Biophys. Res. Commun. 555, 115–120 (2021).

21. C. A. Konopka, S. Y. Bednarek, Variable-angle epifluorescence microscopy: A new way to look at protein dynamics in the plant cell cortex. Plant J. 53, 186–196 (2008).

22. M. Tokunaga, N. Imamoto, K. Sakata-Sogawa, Highly inclined thin illumination enables clear single-molecule imaging in cells. Nat. Methods 5, 159–61 (2008).

23. H. Ueno, et al., Simple dark-field microscopy with nanometer spatial precision and microsecond temporal resolution. Biophys. J. 98, 2014–2023 (2010).

24. T. M. Watanabe, et al., Four-dimensional spatial nanometry of single particles in living cells using polarized quantum rods. Biophys. J. 105, 555–564 (2013).

25. Y. Okada, N. Hirokawa, A processive single-headed motor: Kinesin superfamily protein KIF1A. Science. 283, 1152–1157 (1999).

26. A. Hutterer, M. Glotzer, M. Mishima, Clustering of Centralspindlin Is Essential for Its Accumulation to the Central Spindle and the Midbody. Curr. Biol. 19, 2043–2049 (2009).

27. J. Yajima, M. C. Alonso, R. a Cross, Y. Y. Toyoshima, Direct long-term observation of kinesin processivity at low load. Curr. Biol. 12, 301–6 (2002).

28. E. Berliner, E. C. Young, K. Anderson, H. K. Mahtani, J. Gelles, Failure of a single-headed kinesin to track parallel to microtubule protofilaments. Nature 373, 718–721 (1995).

29. D. Chrétien, S. D. Fuller, Microtubules switch occasionally into unfavorable configurations during elongation. J. Mol. Biol. 298, 663–676 (2000).

30. R. D. Astumian, M. Bier, Fluctuation driven ratchets: Molecular motors. Phys. Rev. Lett. 72, 1766–1769 (1994).

31. M. Kikkawa, N. Hirokawa, High-resolution cryo-EM maps show the nucleotide binding pocket of KIF1A in open and closed conformations. EMBO J. 25, 4187–4194 (2006).

32. L. Cao, et al., The structure of apo-kinesin bound to tubulin links the nucleotide cycle to movement. Nat. Commun. 5 (2014).

33. W. Hwang, M. J. Lang, M. Karplus, Kinesin motility is driven by subdomain dynamics. eLife 6, 1–25 (2017).

34. T. Davies, et al., CYK4 Promotes Antiparallel Microtubule Bundling by Optimizing MKLP1 Neck Conformation. PLoS Biol. 13, 1–26 (2015).

35. S. Can, M. a Dewitt, A. Yildiz, Bidirectional helical motility of cytoplasmic dynein around microtubules. eLife, e03205 (2014).

36. L. Kaplan, A. Ierokomos, P. Chowdary, Z. Bryant, B. Cui, Rotation of endosomes demonstrates coordination of molecular motors during axonal transport. Sci. Adv. 4 (2018).

37. W. Hwang, M. J. Lang, M. Karplus, Force generation in kinesin hinges on cover-neck bundle formation. Structure 16, 62–71 (2008).

38. S. Enoki, et al., High-Speed Angle-Resolved Imaging of a Single Gold Nanorod with Microsecond Temporal Resolution and One-Degree Angle Precision. Anal. Chem. 87, 2079–2086 (2015).

39. M. Castoldi, A. V Popov, Purification of brain tubulin through two cycles of polymerization-depolymerization in a high-molarity buffer. Protein Expr. Purif. 32, 83–8 (2003).

40. S. Ray, E. Meyhöfer, R. A. Milligan, J. Howard, Kinesin follows the microtubule’s protofilament axis. J. Cell Biol. 121, 1083–1093 (1993).

## SI References

1. A. Ramaiya, B. Roy, M. Bugiel, E. Schäffer, Kinesin rotates unidirectionally and generates torque while walking on microtubules. Proc. Natl. Acad. Sci. 114, 10894–10899 (2017).

2. J. Howard, “Mass, Stiffness, and Damping of Proteins” in Mechanics of Motor Proteins and the Cytoskeleton, (Sinauer, 2005), pp. 29–47.

